# Postnatal plasticity in the olfactory system of the juvenile swine brain

**DOI:** 10.1101/2025.08.17.670702

**Authors:** Júlia Freixes, Fatma ElZahraa S. Abdel-Rahman, Roberto Nebbia, Loreta Medina, Ester Desfilis

## Abstract

Swine have an excellent sense of smell and highly complex olfactory brain structures, which play a crucial role in their complex social interactions. In other mammals the olfactory system is known to exhibit significant plasticity, even during adulthood. The aim of this study was to investigate postnatal plasticity in olfactory areas of juvenile swine brains by studying immature cells immunoreactive for the microtubule-associated protein doublecortin (DCX). Using immunofluorescence, we studied DCX coexpression with the cell proliferation marker Ki-67, and different neuronal markers. Our results show the existence of numerous DCX+ cells throughout the olfactory pallial areas. In some of them, we found DCX+/Ki-67+ coexpressing cells, suggesting that they were proliferating. Some of these proliferating cells were grouped in tangentially-oriented migratory-like chains, forming the rostral migratory stream to anterior olfactory area and olfactory bulb. Moreover, chains of DCX+ cells were found in the external capsule and white matter adjacent to the temporal horn of the ventricle. Chains of DCX+ cells were observed crossing the internal layers of the piriform and entorhinal cortices. In layer II of these cortices, DCX+ cells of varying maturity degrees and neuronal phenotypes (including NeuN expression) were present. This suggests the existence of multiple migratory streams along the anteroposterior axis. Most DCX+ immature cells in the migratory chains and in the anterior olfactory area, piriform and entorhinal cortices expressed the transcription factor Brn2 (Pou3f2), suggesting the incorporation of new glutamatergic neurons in these areas. Together, these results highlight the interest of swine to study the role of postnatal brain plasticity and their potential for regeneration in large, gyrencephalic brains.

## INTRODUCTION

Plasticity of the central nervous system refers to its ability to make adaptive changes at functional and structural levels (Bonfanti 2006). It is essential during the development of neural circuits but also plays a crucial role throughout ontogeny for learning, behavioral plasticity, and brain recovery following injury (Kaas 2015). In the juvenile and adult brain of vertebrates, plasticity is related to morphological and/or functional changes of pre-existing neurons (including synaptic plasticity) (Hardy and Saghatelyan 2017), to the continuous production of new neurons (adult neurogenesis) (García-Verdugo et al. 2002; Lledo et al. 2006; Bonfanti and Peretto 2011; Paredes et al. 2015), and to the presence of non-newly born immature neurons (Bonfanti and Nacher 2012; Bonfanti et al. 2024).

In particular, the olfactory systems, from the olfactory epithelia to brain cortical areas, show a great plasticity, involving the incorporation of newly born neurons throughout life in most vertebrates, including mammals (Font et al. 2001; García-Verdugo et al. 2002; Kempermann et al. 2004; Gould 2007; Zupanc 2021). Postnatal neurogenesis has been described in the olfactory bulb (OB) of different mammalian species, including marsupials (Bartkowska et al. 2022) and eutherians such as rodents (Bagley et al. 2007; Brill et al 2009), bats (Chawana et al. 2013), sheep (Brus et al. 2012), humans (Bédard and Parent 2004; Dennis et al. 2016; Sanai et al. 2011) and non-human primates (Kornack and Rakic 2001; Gil-Perotin et al. 2009).

In different species, the cells that incorporate postnatally in the OB are generated in the subventricular zone of the lateral ventricle (SVZ), where a subpopulation of astrocyte-like stem cells produces neuronal progenitors that eventually become neuroblasts (Merkle et al. 2007; Kelsch et al. 2007; Young et al. 2007). After mitosis, these immature cells express doublecortin (DCX), which is crucial for their migration, since this protein regulates the microtubule polymerization and depolymerization (Alvarez-Buylla and García-Verdugo 2002). Immature neuroblasts migrate tangentially forming chains, establishing membrane contacts between them that serve as support for migration, without the guidance of radial glial or axonal fibers (Lois et al. 1996; Alvarez-Buylla and García-Verdugo 2002; Wang et al. 2011). These cellular chains extend from the SVZ along the rostral migratory stream (RMS) to reach the OB, where most of the DCX+ cells mature to produce different subtypes of GABAergic interneurons, expressing calretinin (CR), calbindin (CB) or tyrosine hydroxylase (TH) (Kohwi et al. 2005, 2007; Young et al. 2007). These newly incorporated interneurons establish dendrodendritic connections with mitral and tufted principal neurons of the OB and modulate the activity of these cells for shaping odor representation (Sakamoto et al. 2014). Adult-generated cells constitute a large proportion of neurons in OB and some studies relate them with olfactory learning and discrimination (Barker et al. 2011; Alonso et al. 2012; Li et al. 2018).

DCX+ cells and/or cells expressing other immature cell markers such as PSA-NCAM are also found in other olfactory pallial areas, including piriform and entorhinal cortices and the basal complex of the amygdala, in different mammals, as rodents, sheep and primates (Bernier et al. 2002; Gómez-Climent et al. 2008; Luzzati et al. 2009; Bonfanti and Nacher 2012; Sorrells et al. 2019; Ghibaudi et al. 2023a; Li et al. 2023; Alderman et al. 2024). However, there is a debate about the temporal origin of immature cells in these pallial areas (reviewed in Bonfanti 2006; Bonfanti and Peretto 2011; Jurkowski et al. 2020). Some studies suggest that immature cells found in the postnatal olfactory cortices and pallial amygdala originate from neuroblasts located along the temporal migratory stream (TMS), which extends from posterior parts of the SVZ (Bernier et al. 2002; Cai et al. 2009; Chawana et al. 2013; Ellis et al. 2018; Marlatt et al. 2011; Tonchev et al. 2003). In contrast, based on the apparent absence of mitotic markers in DCX+ cells, it was suggested that these immature neurons likely originate prenatally and remain in a quiescent state until needed, providing a type of non-neurogenic plasticity that is proposed to be more prevalent in the brain of mammals with large gyrencephalic brains (Bonfanti and Nacher 2012; Palazzo et al. 2018; Piumatti et al. 2018; Bonfanti and Couillard-Després 2023; Ghibaudi et al. 2023b, 2025; Bonfanti et al. 2024).

Although the general organization of the olfactory system is similar in all mammalian species, the olfactory brain centers vary substantially in relative size across species (Voogd et al. 1998), in relationship with the importance that the sense of smell plays in the survival and reproduction of each species. Swine are known for their highly developed sense of smell (Moulton 1967). These animals show a high olfactory acuity (Brunjes et al. 2016; Signoret et al. 1975), and they have been employed as a model to reveal the importance of olfactory signals in sexual behaviors (Dorries et al. 1995; Sink 1967), aggressive and submissive behaviors (McGlone 1985; McGlone et al. 1987), and social recognition (Kristensen et al. 2001). Moreover, the swine has recently emerged as an ideal large animal model in biomedical research, especially that related to neuroscience, because its brain is more similar in development, size and structure to the human brain compared to murine models (Lind et al. 2007; Kinder et al. 2019). Furthermore, in the gyrencephalic swine brain the olfactory system is much larger than that of other mammalian species, including other macrosmatic mammals with a lissencephalic brain, such as lab rodents (Brunjes et al. 2016). This makes the swine a good model to study postnatal neurogenesis and other types of plasticity in the olfactory brain.

DCX+ cells have been found in SVZ and RMS of piglets, and their number increases after traumatic cortical injury (Costine et al. 2015). After injections of bromodeoxyuridine (BrdU) in piglets, incorporation of new neurons was shown in the OB and primary olfactory cortex (Martin et al. 2013). Moreover, there is evidence of neuronal precursors and neuroblasts coexpressing the mitotic marker Ki-67 in the ventricular/SVZ of juvenile swine (Torrijos-Saiz et al. 2025). This suggests that, in addition to quiescent immature neurons, there might be some postnatal production of new neurons in swine gyrencephalic brains, at least until juvenile ages. However, the existence of immature neurons and their specific phenotype in olfactory pallial areas of juvenile swine remains unknown.

The aim of the present study was to investigate the plasticity potential throughout the olfactory pallium of juvenile swine (*Sus scrofa domesticus*), including the olfactory bulbs, the anterior olfactory areas, the piriform cortex, the cortical amygdala and the entorhinal cortex. To this aim, we analyzed the expression of DCX, in combination with the cell proliferation marker Ki-67 (Kee et al. 2002), the transcription factor Brn2 that is expressed in subpopulations of pallial glutamatergic neurons (Dominguez et al. 2012; Brunjes and Osterberg 2015), and other neuronal phenotypic markers (CR, TH, NeuN) in different olfactory pallial areas of juvenile 2- to 5-month-old swine of both sexes. To better understand the topological location of the cells with respect to brain subdivisions (for topology concept see Nieuwenhuys 1998) and their potential migratory routes, we analyzed their distribution across different sectioning planes.

## MATERIALS AND METHODS

In the present study, we used brains from swine *(Sus scrofa domesticus*) provided by the Applied Biomedical Research Centre (CREBA) of the Institute of Biomedical Research of Lleida-Dr. Pifarré Foundation (IRBLleida, Spain), located in Lleida (registered as a user center of experimental animals: L9900008; REGA: ES252310036907). In total, we used 6 juvenile swine of both sexes (2 males and 4 females), ranging in age from 2.5 to 3.5 months (, and weighted between 30 to 58kg. At this age, pigs are considered juvenile, as they have not yet reached sexual or full neurodevelopmental maturity. Animals were housed and handled in the pig CREBA facilities, according to the protocol approved by the Animal Research Ethics Committee of the IRBLleida and following the regulations and laws of the European Union (Directive 2010/63/EU) and the Spanish Government (Royal Decrees 53/2013) for the care and handling of animals in research. These animals were previously used for practicing and improving surgical procedures by M.D. surgeons of the Arnau de Vilanova University Hospital of Lleida. After, animals were euthanized, and their brains were extracted and processed as explained below.

### Tissue collection and fixation

For cerebral tissue collection, the animals previously employed for surgery research were sacrificed, just before brain extraction, with a lethal dose of sodium pentobarbital (200mg/kg; IV). After extracting the brain, the tissue was washed with abundant water to remove blood and hemispheres were separated and fixed by immersion in a solution of 4% paraformaldehyde (PFA) in 0.1 M phosphate buffer (PB), for 2-4 days.

### Sample preparation and sectioning

Following fixation, the tissue was cryoprotected in a solution of increasing glycerol concentrations (10% and 20%) and 2% DMSO in 0.1M PB, for 6-10 days. Some hemispheres were cut into four blocks (frontal, temporoparietal, occipital and cerebellum), while others were conserved in a single piece. The tissue was frozen by immersion in -65/-70°C isopentane (2-methyl butane, Sigma-Aldrich, Germany) for 1-2min, following the protocol of Rosene et al. (1986). Frozen brains were wrapped in aluminum foil and stored at -80°C until further use.

For sectioning, samples were embedded in 30% sucrose in 0.1M PB and cut using a freezing sliding microtome (Microm HM 450; Thermo Fisher Scientific), equipped with a large size Freezing Stage (BFS-40MPA, Physitemp Instruments, LLC, USA). Coronal, sagittal and horizontal sections of 100μm thick were collected at 4°C in 0.1M PB, into twelve parallel series with about 20 sections each block for coronal sections and each sagittal- or horizontally sectioned hemisphere. For each series, selected sections of the appropriate levels including the areas of interest were processed for studying. The sections not used immediately were stored in the 30% sucrose solution and frozen at -20°C.

In one animal (male), each brain hemisphere was sectioned in a different plane (coronal and sagittal), while in the other animals (four females and one male), only one brain hemisphere was sectioned, obtaining one coronal (female), one sagittal (female) and three horizontal (two females and one male) section series. Regarding the coronal sections, four antero-posterior levels of the telencephalon were studied (Fig. 1A). The most anterior level (Fig. 2A) was equivalent to 36.375mm and the most posterior to -1.125 mm antero-posterior stereotaxic coordinates from the MRI Atlas (Saikali et al. 2010). Overall, 20 sections were Nissl-stained, about 230 sections were processed for immunohistochemistry and 150 sections for double or triple immunofluorescence.

**Figure 1.**
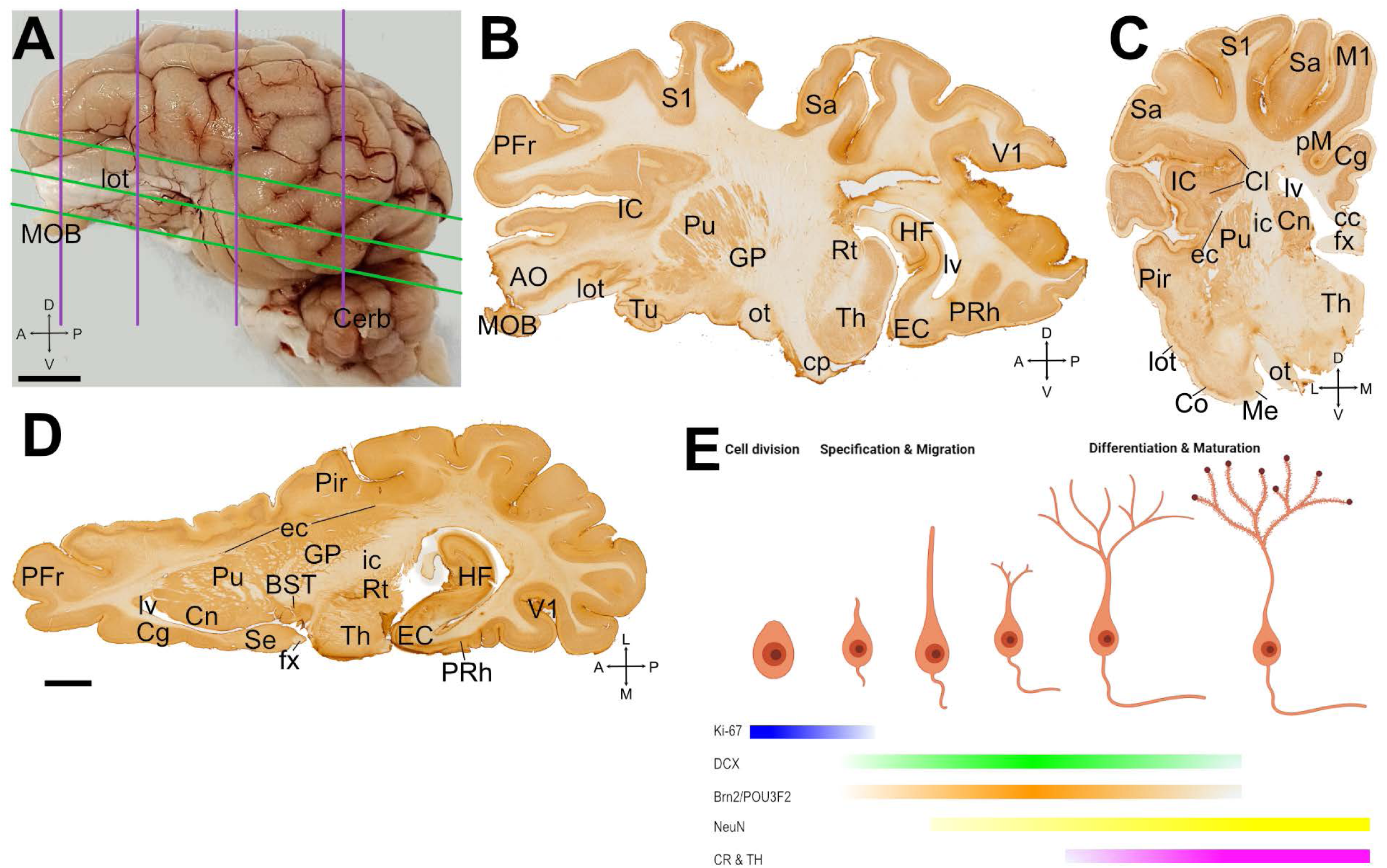
Swine brain, section planes and temporal sequence of expression of cellular markers used in this study. (A) Lateral view of the left hemisphere of swine brain with lines representing the planes used for obtaining frontal (purple) and horizontal (green) sections. (B-D) Examples of sagittal (B), coronal (C) and horizontal (D) sections immunostained for NeuN. Anteroposterior (A-P), dorsoventral (D-V) and mediolateral (M-L) axes are provided in A-D for orientation. (E) Diagram of the developmental sequence of newborn neurons showing the morphological and functional stages in relation to the temporal expression pattern of cellular markers used in this study. For abbreviations, see list. Scales: A = 1mm, D = 5mm (applies to B-D)

**Figure 2.**
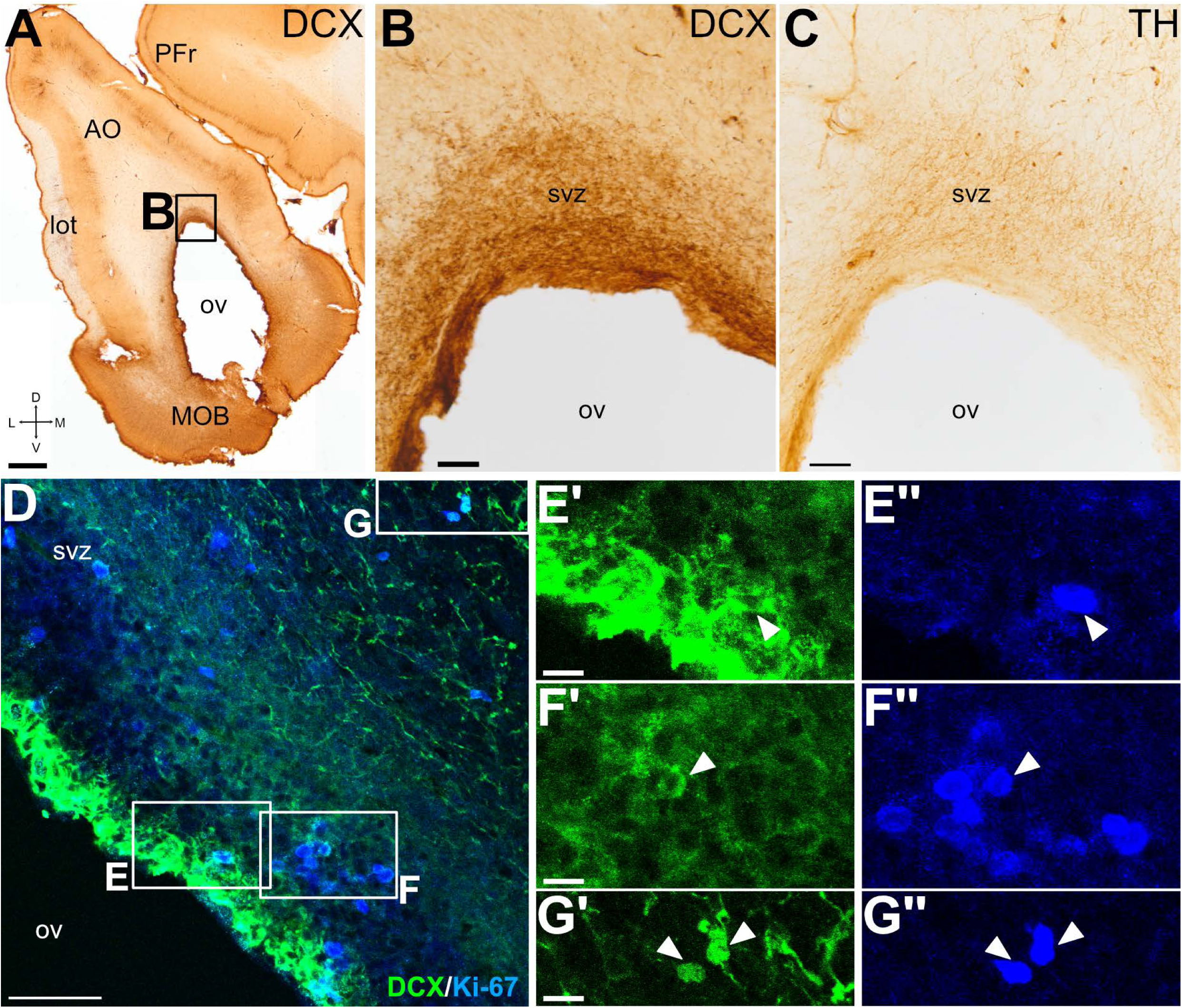
Proliferative capacity of the DCX+ cells in the SVZ around the rostral extension of the lateral ventricle. (A-G) Frontal sections at similar level to allow visualization of the olfactory ventricle (ov), as well as the main olfactory bulb (MOB) and anterior olfactory area (AO), immunohistochemically stained for DCX (A-B) and TH (C), or double immunofluorescence stained (D-G) for DCX (green) and Ki-67 (blue). Squared area in A is shown at higher magnification in B, while squared areas in D are shown at higher magnification in E’-E’’, F’-F’’ and G’-G’’. Arrowheads point to examples of cells coexpressing DCX and Ki-67. These are more abundant in SVZ close to the ependymal layer, but some are also observed external layer of SVZ, which is part of the rostral migratory stream (RMS). Mediolateral and dorsoventral axes are indicated in A for orientation. For other abbreviations, see list. Scales: A = 1mm; B, C = 100µm; D = 50µm; E’, F’, G’ = 10µm (also applies to E’’, F’’, G’’)

### Nissl staining

To study the tissue organization and to help with the identification of the areas, some sections were processed for Nissl staining. After being mounted with a solution of gelatine from porcine skin at 0.5% in Tris buffer and dried, sections were incubated for 1 minute with filtered Toluidine blue 1% in acetic buffer (composed of acetic acid 0.2 M and sodium acetate 0.2 M) at pH 4.6.Sections were then rinsed with a solution of acetic acid 0.01% in ethanol 70%, to remove excess of staining. Subsequently, the slides were dehydrated with ascending ethanol concentrations (70%, 96% and 100%), followed by a brief xylene wash, and finally coverslipped with Permount mounting medium (Thermo Fischer Scientific).

### Immunohistochemistry

Free-floating brain sections were processed for immunohistochemistry to detect different markers: Ki-67, doublecortin (DCX), calretinin (CR), tyrosine hydroxylase (TH), neuronal nuclei (NeuN) and Pou3f2/Brn2 TF (Brn2), using specific antibodies (Table 1). Sections were washed with 0.1M PB and then processed for antigen retrieval by incubation in a sodium citrate buffer (10mM Sodium Citrate, 0.05% Tween 20, pH 6.0)for 1h at 60°C. After rinsing, sections were permeabilized with PB containing 0.3% Triton-X 100 (PB-Tx; 0.1 M), and incubated for 2h at room temperature in a blocking solution containing 2% of bovine serum albumin (BSA) and 20% of normal rabbit, horse or goat serum (according to the species in which the secondary antibody was raised) in 0.1M PB. Next, sections were incubated in primary antibody solution (Table 1) in PB-Tx for 48h-72h at 4°C with gentle shaking. After rinsing, sections were incubated in a biotinylated secondary antibody (Table 2) in PB-Tx overnight, at 4°C with gentle shaking. Successively, sections were washed, and endogen peroxidase was blocked with a solution of PB with 20% methanol and 2% H_2_O_2_ for 20min. After, sections were washed and incubated in the avidin–biotin complex (AB Complex, Vector Laboratories Ltd.) for 1 h at room temperature. After incubation in the ABC solution, the sections were washed with one rinsing of PB, followed by two of Tris buffer (0.05M, pH 7.6).

**Table 1.**
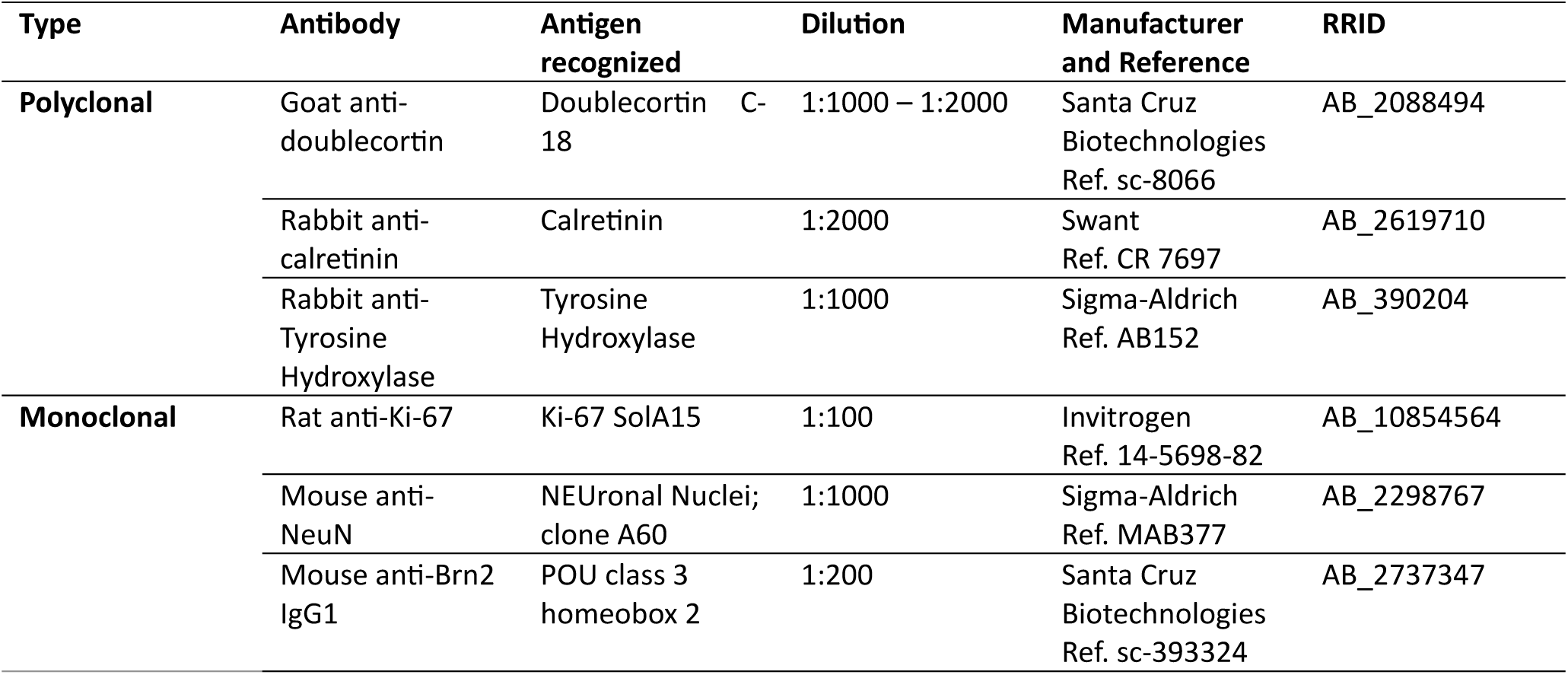
Primary antibodies.

**Table 2.** Secondary antibodies.

| Type | Antibody | Dilution | Manufacturer and Reference | RRID |
| --- | --- | --- | --- | --- |
| **Biotinylated** | Rabbit anti-goat IgG (H+L), Biotinylated | 1:200 | Vector Laboratories<br>Ref. BA-5000 | AB_2336126 |
|  | Goat anti-mouse IgG (H+L), biotinylated | 1:200 | Vector Laboratories<br>Ref. BA-9200 | AB_2336171 |
|  | Goat anti-rabbit IgG (H+L), Biotinylated | 1:200 | Vector Laboratories<br>Ref. BA-1000 | AB_2313606 |
| **Fluorescent** | Donkey anti-goat IgG (H+L), coupled to Alexa 488 | 1:500 | Jackson ImmunoResearch<br>Ref. 705-546-147 | AB_2340430 |
|  | Donkey anti-mouse IgG (H+L), coupled to Alexa 488 | 1:500 | Invitrogen<br>Ref. A21202 | AB_141607 |
|  | Donkey anti-goat IgG (H+L), coupled to Alexa 568 | 1:500 | Invitrogen<br>Ref. A11057 | AB_2534104 |
|  | Donkey anti-rabbit IgG (H+L), coupled to Alexa 568 | 1:500 | Invitrogen<br>Ref. A10042 | AB_2534017 |
|  | Goat anti-rabbit IgG (H+L), coupled to Alexa Plus 405 | 1:250 | Invitrogen<br>Ref. A48254 | AB_2890548 |
|  | Goat anti-mouse IgG (H+L), coupled to Alexa Plus 405 | 1:250 | Invitrogen<br>Ref. A48255 | AB_2890536 |
|  | Goat anti-rat IgG (H+L), coupled to Alexa Plus 405 | 1:250 | Invitrogen<br>Ref. A48261 | AB_2890550 |

Following this, the sections were incubated with diaminobenzidine (DAB, SIGMAFAST™ tablets, Sigma-Aldrich Co. LLC), containing urea and H_2_O_2_, diluted in MilliQ water according to the manufacturer’s instructions. The reaction was stopped by rinsing the sections with Tris buffer. Sections were then mounted, dehydrated and covered with Permount™ as described before.

### Double and triple immunofluorescence

Multiple labeling immunofluorescence was performed to detect cells coexpressing different markers. Following tissue permeabilization and blocking of non-specific binding, sections were incubated at 4°C for 48h-72h with a cocktail of primary antibodies (Table 1). After rinsing, sections were incubated overnight at 4°C in PB-Tx with the corresponding cocktail of fluorescent secondary antibodies (Table 2). Finally, the sections were mounted with a solution of gelatine at 0.5% and covered with an antifading mounting medium (Vectashield Hardset Antifade mounting medium, Vector Laboratories Ltd.).

### Negative controls for immunohistochemistry and immunofluorescence

Horizontal and sagittal sections at the level of anterior olfactory area (AO), piriform cortex (Pir), entorhinal cortex (EC), and main olfactory bulb (MOB) were processed following all the steps for immunohistochemistry or triple immunofluorescence but omitting primary antibodies. No specific cell labeling was observed (Supplementary figure).

### Identification of regions of interest

To identify the regions of interest, we used the brain atlas of the wild boar in Nissl staining from the University of Wisconsin-Madison, as well as the MRI 3D atlas from Saikali et al. (2010). Our identification was based on the topological position of the structures plus our Nissl staining and NeuN immunostained parallel sections.

### Digital photographs and figures

Digital microphotographs of the sections processed for immunohistochemistry were taken on a Leica microscope (DMR HC and DM2500 LED, Leica Microsystems GmbH) equipped with a Zeiss Axiovision Digital Camera (Carl Zeiss). Serial images from fluorescent material were taken under confocal microscope (Olympus FV1000; Olympus Corporation). Selected digital fluorescent images were adjusted and extracted using Olympus FV10-ASW 4.2 Viewer (Olympus Corporation). Finally, the figures were mounted using Affinity Designer 2 (version 2.5.3, Serif Corporation).

## RESULTS

We analyzed coronal, sagittal and horizontal sections at different pallial levels from juvenile swine brains of both sexes to carry out a systematic description of the distribution of DCX+ cells throughout the subdivisions of the olfactory pallium (Fig. 1A-D). The expression pattern of DCX in olfactory pallial areas was similar across animals and in both sexes. However, since we had a limited brain number, our results did not allow a reliable comparison between sexes or ages. DCX+ cells were found throughout all the olfactory pallial regions analyzed, including the OB, the anterior olfactory areas (AO), the piriform cortex (Pir), the cortical areas of the amygdala (Co) and the entorhinal cortex (EC). As many of the DCX+ cells had a migratory cell morphology and formed chains (in agreement with Torrijos-Saiz et al. 2025), we also explored the possible migratory pathways these immature cells might follow from progenitor areas to their final position. We studied the expression of Ki-67, a mitotic marker to identify proliferative zones. In addition to know the phenotype of DCX+ cells we carried double immunofluorescence using Brn2 (Pou3f2) (which is expressed in subpopulations of immature glutamatergic cortical neurons), NeuN (that is expressed in neurons since they start to differentiate) and other markers of mature neurons (TH, CR). The temporal expression of these different markers and DCX, from cell proliferation till maturation, is represented in Fig. 1E, pointing to overlapping windows for exploring possible coexpression in immature cells.

### Postnatal proliferative potential around the rostral part of the lateral ventricle and RMS

In swine, the lateral ventricle (lv) had a large extension, both rostral- and caudally, and included a temporal horn (Fig. 1). The rostral part of the lv extended through the AO and reached the OB (this most rostral extension of the ventricle is named olfactory ventricle, ov), showing abundant DCX+ somas and fibers surrounding the wall of the ventricle (SVZ, Fig. 2A,B,D), particularly in its dorsal part (Fig. 2B). In the same region, we also saw TH+ fibers, some of which reached the ependymal layer (Fig. 2C).

To know if these immature DCX+ cells were generated postnatally, we studied the expression of the cell-division marker Ki-67. Ki-67+ somas were abundant through the ventricular zone (V) and, especially, in the SVZ (Fig.2D) and some of those in SVZ coexpressed DCX (details in Fig. 2E’-F’’ from squared areas in Fig. 2D). We also found double-labeled cells coexpressing Ki-67 and DCX with a farther location in the most external layer of SVZ (detail in Fig. 2G’-G’’), identified as layer 3 by Torrijos-Saiz et al. (2025), which is part of RMS. The coexpression of Ki-67 and DCX in cells located at the SVZ of the olfactory ventricle indicates that they proliferate *in situ* within the OB/AO.

In addition, in sagittal (Fig. 3A) and horizontal sections, we observed numerous DCX+ cells with an elongated morphology, forming large tangentially oriented chains, with a posteroanterior alignment, resembling the RMS (arrowheads in Fig. 3B-C). Some of these cells with an elongated migratory-like morphology (arrowheads in Fig. 3D) showed coexpression of Ki-67 (Fig. 3E-E’’), suggesting that they are dividing during migration, and some of them were oriented parallel to TH+ fibers and to blood vessels (Fig. 3F-F’’). Moreover, at AO levels, many of the DCX+ cells in the chains showed coexpression of the transcription factor Brn2 (Fig. 3G-I’’’). This was especially evident for the DCX+ cells of RMS in the dorsal part of AO (i.e. above ov; Fig. 3H-H’’’), but some of those in the ventral side also coexpressed Brn2 (Fig. 3I-I’’’).

**Figure 3.**
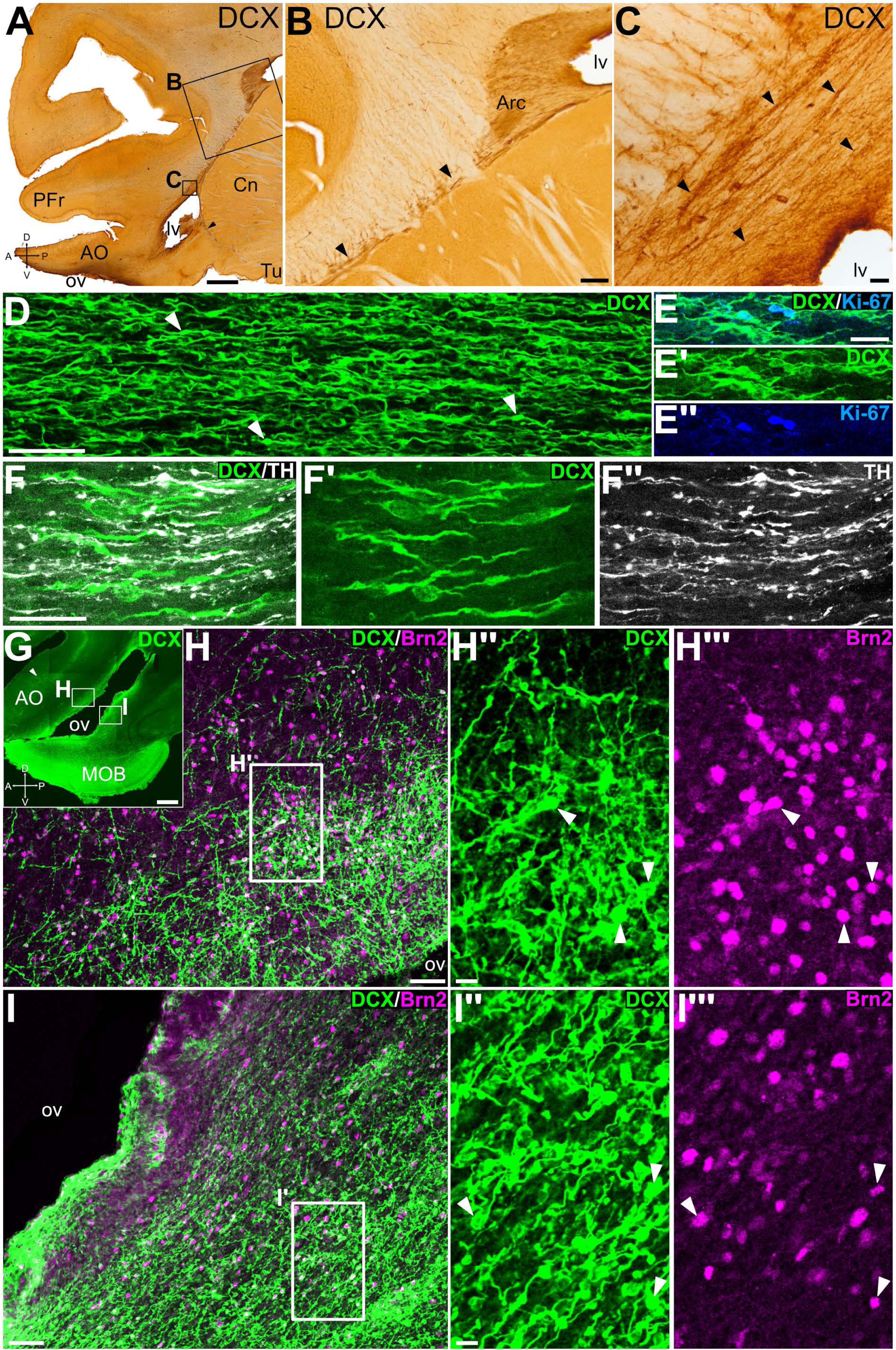
Proliferation and phenotype of DCX+ cells of the rostral migratory stream. Sagittal sections (panoramic views in A and G; details in the rest) immunohistochemically stained for DCX (A-C), or immunofluorescence stained (D-I’’’) for DCX (green), Ki-67 (blue), TH (white) and/or Brn2 (magenta). Anteroposterior (A-P) and dorsoventral (D-V) axes are provided in some of the images for orientation. Squared areas in A are shown at higher magnification in B and C; squared areas in G are shown at higher magnification in H and I; and squared areas in H and I are shown in H’’-H’’’ and I’’-I’’’, respectively. (B) Detail of the SVZ around the lateral ventricle, showing abundant DCX+ cell patches located in the Arc, as well as chains of elongated DCX+ cells that extend rostrally from this region along the rostral migratory stream (RMS, arrowheads). (C) Detail of the tangentially oriented DCX+ cellular chains (arrowheads) in the RMS. (D) Confocal detail of the chains of DCX+ elongated cells (arrowheads point to some examples) in the RMS, in a position similar to that of the chains in C. (E-E’’) Detail of the RMS from another sagittal section double labelled for DCX and Ki-67 (at a similar level to that in C), showing chains with DCX+ elongated cells, some of which coexpress Ki-67. (F-F’’) Confocal images of a detail of the RMS (level similar to C), showing DCX+ cells with a parallel orientation to TH+ fibers. (H-I) Confocal images of details of the sagittal section seen in (G) double labeled for DCX and Brn2, showing the dorsal (H) and ventral (I) subdivisions of the olfactory area (OA) around the olfactory ventricle (ov). (H’’-H’’’) Detail of the dorsal AO, showing DCX+ cells that are grouped forming chains with multiple orientations. Some of these DCX+ cells co-express Brn2 (arrowheads). (I’’-I’’’) Detail of the ventral AO, showing DCX+ cells that are grouped forming chains which are oriented parallel to the wall of the ventricle. Some of these DCX+ cells co-express Brn2 (arrowheads). For other abbreviations, see list. Scales: A = 2mm; B = 100µm; C, D, H, I = 50µm; E = 25µm (applies to E-E’’); F = 25µm (applies to F-F’’); G = 1mm; H’’, I’’ = 10µm (applies to H’’, H’’’, I’’, I’’’)

### Distribution and phenotype of DCX+ cells in the olfactory bulbs

We studied the distribution of these immature cells throughout the layers of the main OB (MOB) (Fig. 4). In the glomerular layer (GL), the external plexiform layer (EPL) and mitral cell layer (MCL), the expression of DCX was mostly found in processes but extremely few somas were also seen in the GL and EPL (Fig. 4B-D). Almost all cells expressing DCX in MOB were located in the granule cell layer (GCL) and they were surrounded by DCX+ neuropile; many of the DCX+ cells had a small soma and displayed radially oriented processes, perpendicular to the layers of the OB, which extended to the MCL and beyond (Fig. 4B-D).

**Figure 4.**
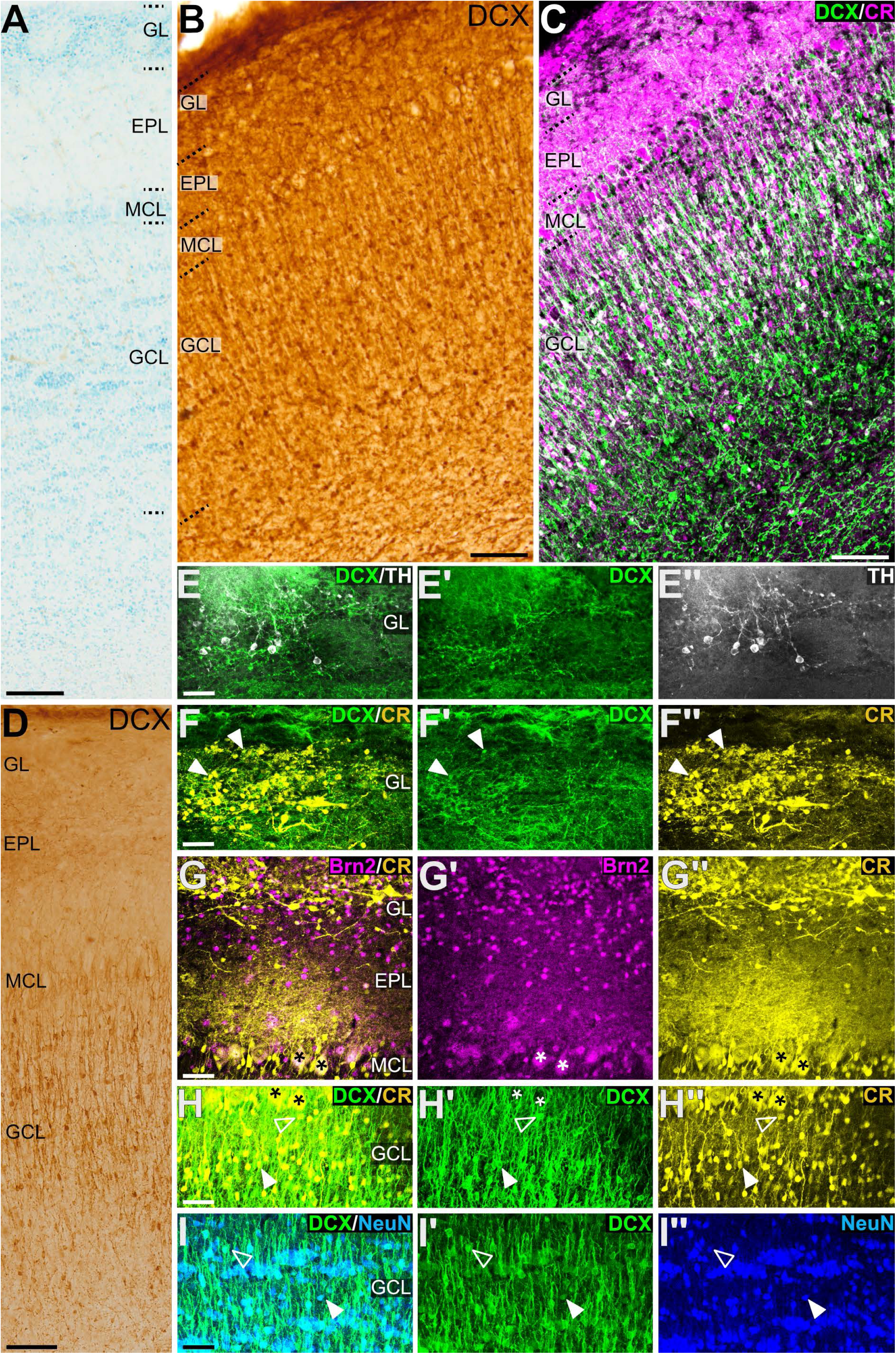
DCX expression pattern in the main olfactory bulb and phenotype of DCX+ cells. (A-D) Details of the main olfactory bulb (MOB) in horizontal sections (A-C, like that shown in Fig. 5A) or sagittal sections (D, like that in Fig. 1B) stained for Nissl (A), immunohistochemistry for DCX (B, D) or double immunofluorescence for DCX (green) and CR (magenta) (C). Layers of the MOB are indicated: glomerular layer (GL), external plexiform layer (EPL), mitral cell layer (MCL) and granular cell layer (GCL). (E-I’’) Confocal images showing details of MOB in sagittal sections at higher magnification, stained by double immunofluorescence for DCX (green), TH (white), CR (yellow) and/or NeuN (blue). (E-E’’) Detail of DCX+ neuropile and TH+ periglomerular cells in GL. (F-F’’) Detail of CR+ periglomerular cells pointing to some examples of coexpression with DCX (arroheads). (G-G’’) Detail of CR+ and/or Brn2+ cells in the MCL, EPL and GL. Coexpression is seen in mitral cells (asterisks) and in some cells of the EPL. (H-H’’) Detail of double-labeled cells in the GCL co-expressing DCX and CR, and mitral cells (asterisks) expressing CR and surrounded by DCX+ apical processes from GCL cells. (I-I’’) Detail showing DCX+ cells co-expressing the neuronal marker NeuN in the GCL. Empty arrowheads in H-I’’ show cells with high signal of DCX and weak signal of CR or NeuN (possibly at an early maturation stage), while filled arrowheads point to cells with weak signal of DCX and high expression of neuronal phenotypes (possibly at a later maturation stage). For other abbreviations, see list. Scales: A, B, C, D = 100µm; E, F, G, H, I = 50µm (applies to E-E’’,F-F’’,G-G’’,H-H’’,I-I’’)

To explore the phenotype of these DCX+ cells, we carried out double immunofluorescence for DCX and either TH, CR, Brn2, or NeuN (Fig. 4E-I’’). We found TH+ periglomerular somas in the GL but none of them coexpressed DCX (Fig. 4E-E’’). We also observed CR+ periglomerular cells, some of which appeared to coexpress DCX (Fig. 4F-F’’). In the MCL, the glutamatergic mitral cells did not express DCX, but they expressed CR and Brn2 (asterisks in Fig. 4G-G’’). We observed a dense DCX+ neuropile surrounding the glutamatergic mitral cells (Fig. 4H-H’’). Most somas expressing DCX in the GCL exhibited a neuronal phenotype, as they expressed NeuN (Fig. 4I-I’’) as well as CR (Fig. 4H-H’’). In both phenotypic markers (NeuN and CR) we found differences in intensities of expression related to differences in DCX immunolabeling, but with opposing gradients: some neurons showed high signal of DCX and weak signal of NeuN or CR (Fig. 4H-I’’, empty arrowheads), while others showed the opposed pattern (Fig. 4H-I’’, white arrowheads). This variability in expression intensity may reflect differences in the maturation degree of these neurons as they transition from an immature to a mature state.

Regarding the accessory olfactory bulb (AOB), it is much smaller than the MOB and can be identified posteromedial to the MOB in horizontal sections (Fig. 5A,B), as previously described in swine (Brunjes et al., 2016). We found some small-sized DCX+ cells, which may correspond to granular cells.

**Figure 5.**
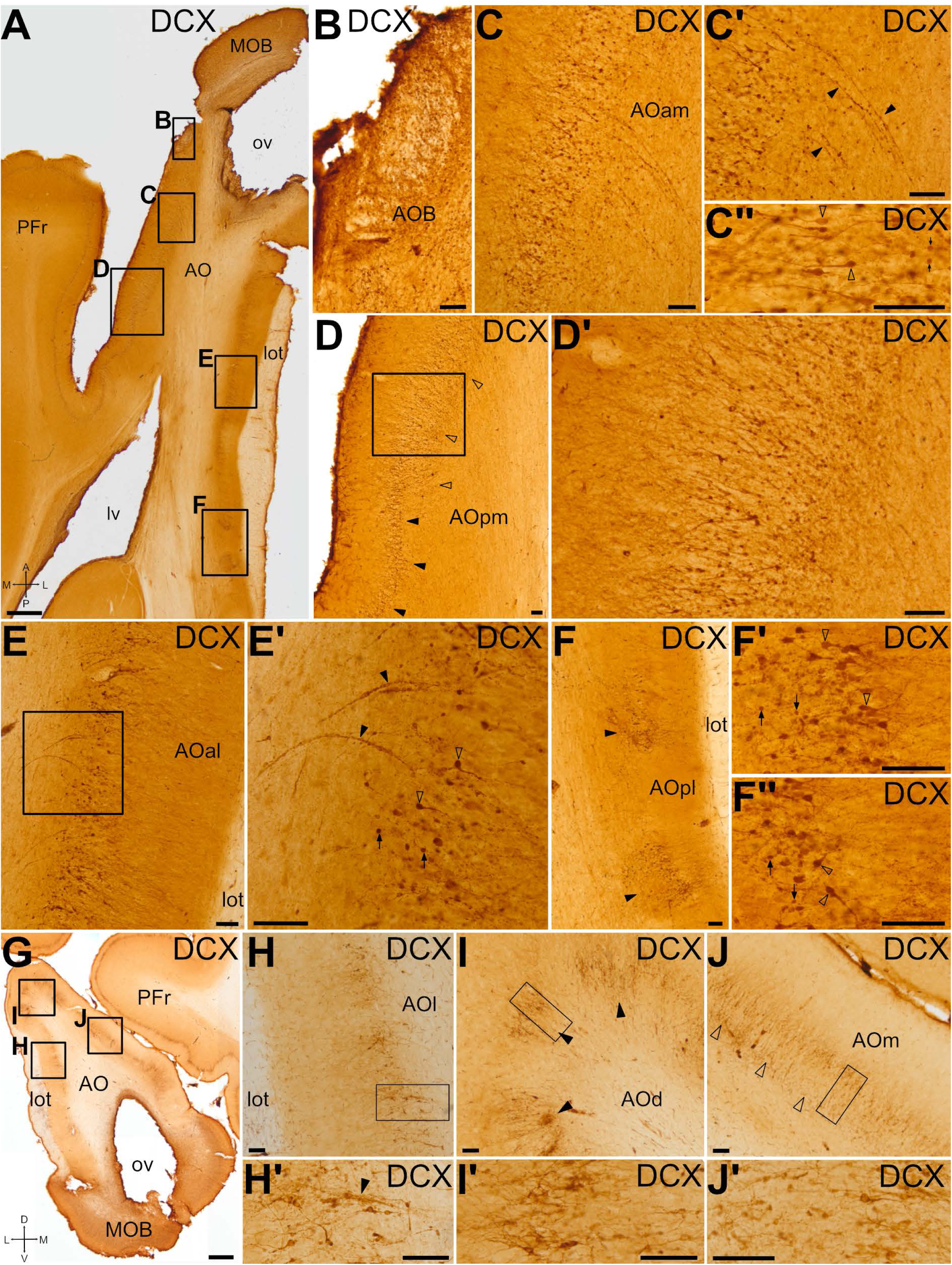
DCX expression pattern in the anterior olfactory area. Horizontal (A-F’’) and frontal (G-J’) sections at the level of the anterior olfactory area (AO) and the main and accessory olfactory bulbs (MOB, AOB), immunohistochemically stained for DCX. (B) shows a detail of the AOB. (C-F’’) shows details in different subdivisions of AO. (C) Detail of DCX+ cells in the anteromedial AO (AOam). (C’) Detail of chains of small-rounded somas expressing DCX in AOam (black arrowheads), extending from the inner part to the cell layer. (C’’) Detail of large cells in AOam (empty arrowheads), with a clear apical dendrite, and small rounded cells (black arrows). (D) Detail of the posteromedial part of the AO (AOpm), showing a thick layer of DCX+ cells (empty arrowheads), which transition to a narrower layer caudally (black arrowheads). (D’) Detail of squared area in D, showing DCX+ cells with small somas without or with simple arborizations in the inner part of the cell layer, and larger somas and more complex dendrites in the external part. (E) Detail of the DCX+ cells in the anterolateral part of the AO (AOal), adjacent to the lateral olfactory tract (lot). (E’) Detail of the squared area in E. As in the AOam, different cell morphologies were identified, including large cells with a clear apical dendrite (empty arrowheads) as well as small-rounded somas without visible arborizations (black arrows). Moreover, chains of DCX+ small-rounded somas were observed (black arrowheads). (F) Detail of posterolateral AO (AOpl), showing scattered patches of DCX+ cells (arrowheads). (F’-F’’) Details of DCX+ cells in AOpl, including large cells with clear dendrites (empty arrowheads) as well as small-rounded somas without visible arborizations (black arrows). (H-J) Details of lateral (H), dorsal (I) and medial (J) parts of anterior AO from the coronal section shown in (G), showing the distribution of DCX+ cells. Details of the cells are shown in H’-J’ (from the squared areas in H-J). The detail in (H’) shows DCX+ cells and a cellular chain (arrowhead) of small DCX+ somas in lateral AO (AOl). (I-I’’) Detail of dorsal AO (AOd), showing patches of DCX+ cells (black arrowheads). (J-J’) DCX+ cells in medial AO (Aom) organized in a double-layer (empty arrowheads), with small somas in the inner part and more complex larger cells in the external part. For other abbreviations, see list. Scales: A, G = 1mm; B-F’’, H’-J’ = 100µm

### Distribution and phenotype of DCX+ cells in the anterior olfactory areas

After leaving the OB, the anterior olfactory area (AO) is the first major target of the mitral and tufted cells projections (López-Mascaraque et al. 1996). In swine, we identified the AO posterior to MOB and AOB and it had a very large extension (as described in Brunjes et al. 2005). The lateral olfactory tract (lot) was observed on AO lateral side, along its entire anteroposterior extension. The swine AO shows a laminar (cortical) structure and, based on differences in lamination, we distinguished at least dorsal, ventral, lateral, medial, and posterior subdivisions. Regarding the DCX expression pattern, we observed a laminar distribution of DCX+ cells in cortical layer II (the cellular layer), with differences depending on the subdivision (Fig. 5A,G). These differences were more evident in the anterior AO, with a trend for a single layer of DCX+ cells in lateral subdivision (Fig. 5E,H, details in E’ and H’), which transitioned to a patchy distribution of cells dorsally (Fig. 5I black arrowheads; detail in I’), and then to a double layer of DCX+ cells in the medial subdivision (Fig. 5C, J detail in J’). The posterior subdivision of the AO (AOp) showed transitional patterns in the distribution of DCX+ cells toward those found in the piriform cortex (laterally) or the prefrontal cortex (medially). The lateral part of the posterior subdivision (AOpl) included distinct clusters of DCX+ cells (Fig. 5F, black arrowheads, details in F’ and F’’). In the medial part of the AOp (AOpm), DCX+ cells were distributed forming a thick band, occasionally appearing as a double layer, parallel to the surface (empty arrowheads in Fig. 5D; detail in D’). This pattern gradually transitioned to a thinner and more organized layer at more posterior levels (black arrowheads in Fig. 5D), until DCX+ cells were restricted to cortical layer II of the prefrontal cortex (Fig. 5A).

In all subdivisions, we were able to distinguish DCX+ neuroblasts with small somas, either lacking or bearing simple arborizations located in the inner part of the DCX+ cell layer, while DCX+ cells with bigger somas and more complex apical dendrites (oriented perpendicular to the brain surface) were found in the outer part of the cell layer (Fig. 5C’’,D’,E’,F’,F’’,H’,I’,J’), resembling the subtypes previously described based on Golgi staining (Brunjes et al., 2016). In the inner part of the AO, we also observed many bead-like chains of DCX+ cells (black arrowheads in Fig. 5C’,E’).

Double immunofluorescence revealed Ki-67 expression in some DCX+ cells located in the inner part of the AO (Ki-67, Fig. 6A, details in A’, A’’), and some small neurons coexpressing DCX and CR (Fig. 6B-C’’). However, other CR+ neurons in the same area did not coexpress DCX. In addition to the strong expression of CR in the axons of the lot, we detected abundant CR+ varicose fibers in close proximity to DCX+ cells in the AO. Some of these fibers ran obliquely and appeared to diverge from the lot (Fig. 6B, detail of the CR+ axons in D’’). In the dorsal AOp, most DCX+ cells coexpressed Brn2 (Fig. 6F,J-, white arrowheads in G’-G’’), and at least a subset also coexpressed CR (empty arrowheads in Fig. 6J-J’’’). Deeper to this DCX+ cell layer, small cells grouped in chains coexpressed DCX and Brn2 (empty arrowheads in Fig. 6H’-H’’, I’-I’’).

**Figure 6.**
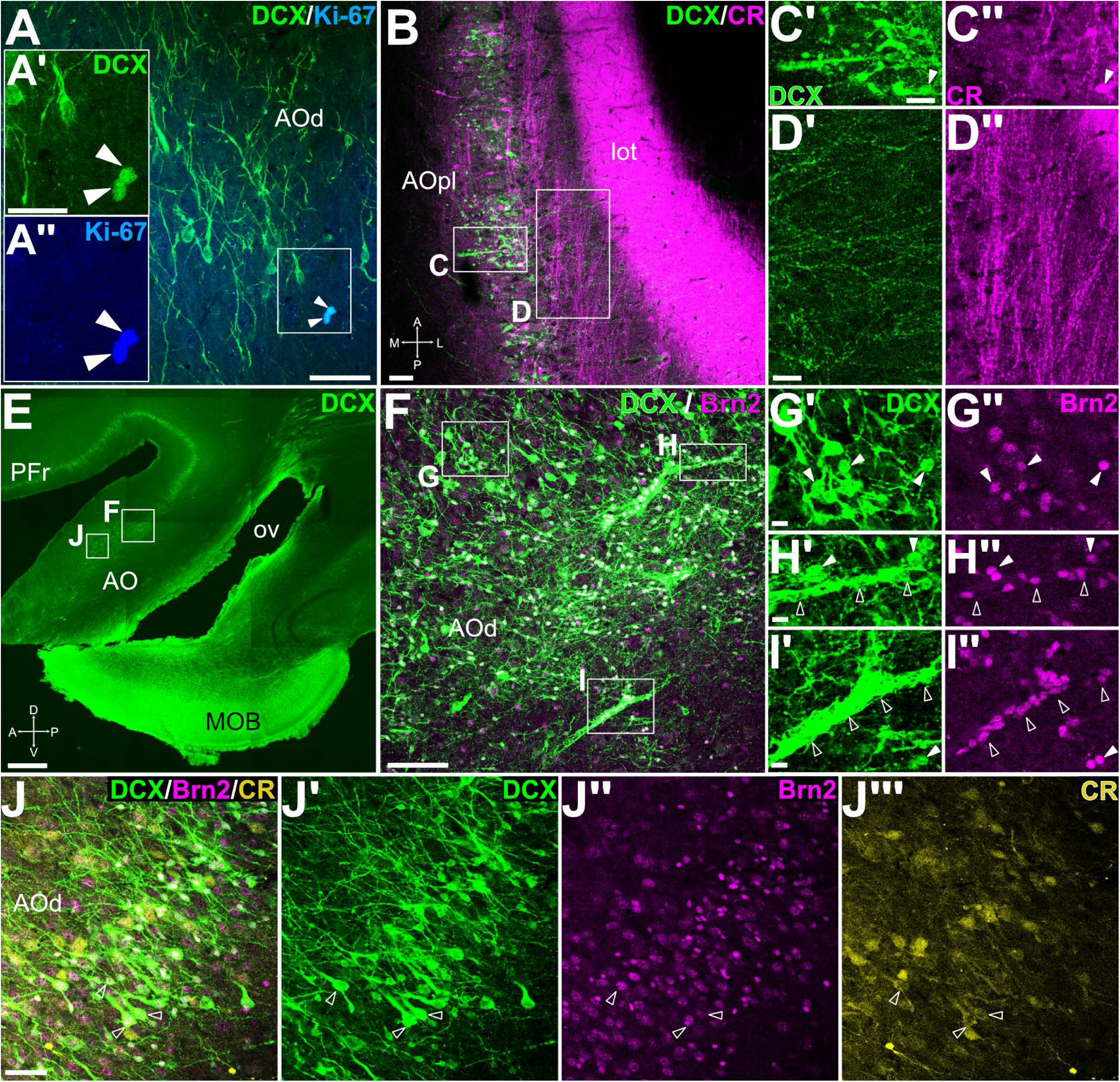
Phenotypes of DCX+ cells in the anterior olfactory area. (A) Confocal image of a frontal section (similar level to Fig. 5G) immunofluorescence stained for DCX (green) and Ki-67 (blue), showing double labeled cells in the dorsal part of the anterior olfactory area (AOd) (arrowheads in the squared area; separate confocal channels shown at higher magnification in A’-A’’). (B) Confocal image of a horizontal section (similar level at Fig. 5A) processed for double labeling immunofluorescence for DCX (green) and CR (magenta), at the level of posterolateral AO (AOpl). Note the strong CR immunoreaction in lateral olfactory tract (lot), and the axons departing from it to approach the DCX+ cells in AO. (C-C’’) Detail showing some immature neurons co-expressing DCX and CR (arrowheads in C’ and C’’). (D-D’’) Detail showing CR+ axons separating from the lot, entering the AOpl, and approaching the apical dendrites of DCX+ cells. (E-J’’’) Confocal images of a sagittal section (E; same as Fig. 3G) and details of dorsal AO (AOd) (F and J, from squared areas in E), processed by immunofluorescence for DCX (green), Brn2 (magenta) and CR (yellow). (F) Detail of cells showing co-expression of DCX and Brn2. (G-G’’) detail of the external part of the AOd cell layer, showing double labeled cells (arrowheads). (H-I’’) Details of the inner part of the AOd cell layer, showing double labeled cell chains (empty arrowheads), as well as coexpressing individual cells with small-rounded somas (filled arrowheads). (J-J’’’) Detail of cells co-expressing DCX, Brn2 and CR, with a high intensity of DCX and a weaker intensity of Brn2 and CR (empty arrows). For other abbreviations, see list. Scales: A = 50µm; A’= 25 µm (also applies to A’’); B = 100µm; C’, D’ = 50µm (also applies to C’’, D’’). E = 1mm; F = 100µm; G’, H’, I’ = 10µm (also applies to G’’, H’’ and I’’); J = 50µm (applies to J-J’’’)

### Proliferation, migration and integration of immature cells in the piriform cortex

The piriform cortex (Pir) is one of the main recipients of the OB mitral and tufted cells projections through the lot (Martínez-Marcos 2009; Shipley et al. 2008; Martínez-García et al. 2012). In swine’s brain, the Pir can be followed from the AO until lateral entorhinal cortical levels, parallel to the lot, (Figs. 1, 7A). As noted above, the axons of mitral/tufted cells travelling in the lot are immunoreactive for CR, and this is a useful marker to reliably locate the Pir in different section planes (Fig. 7B,E). The main, thickest part of the lot is adjacent to the ventral Pir from anterior to posterior levels (Figs. 1A; 7A,D,E). As in other mammals, the Pir has three layers with different cell density, as evidenced with Nissl staining: layer I is the most superficial and is a plexiform layer with very few cells, layer II has a high neuron density, and layer III is the deepest and shows a moderate cell density (Fig. 7D). Calretinin immunostainings allowed to subdivide layer I of Pir in a superficial sublayer Ia, with densely packed CR+ fibers, and a deep sublayer Ib with apical, spiny dendrites of layer II neurons (Fig. 7B,E) and CR+ varicose fibers running mostly parallel to the surface (asterisks in Fig. 7B). Abundant DCX+ cells with different morphologies were mainly found in layer II from rostral to caudal levels of the Pir (Fig. 7C,F). Many DCX+ cells displayed a pyramidal or pear-shaped soma, relatively large, located in cortical layer II and a complex dendritic arborization, with a radially oriented apical dendrite directed toward the lot and some basal dendrites, resembling the pyramidal cells described in the piriform cortex of other species (black arrowheads in Fig. 7G,H). The apical dendrite of these cells extended to the external part of layer I (layer Ia), that receives the CR+ olfactory input from MOB. Moreover, in the external part of layer II we found DCX+ round or crescent-shaped cell bodies with two or more prominent apical dendrites, resembling semilunar cells described previously in other mammals (white arrowheads in Fig. 7G,H).

**Figure 7.**
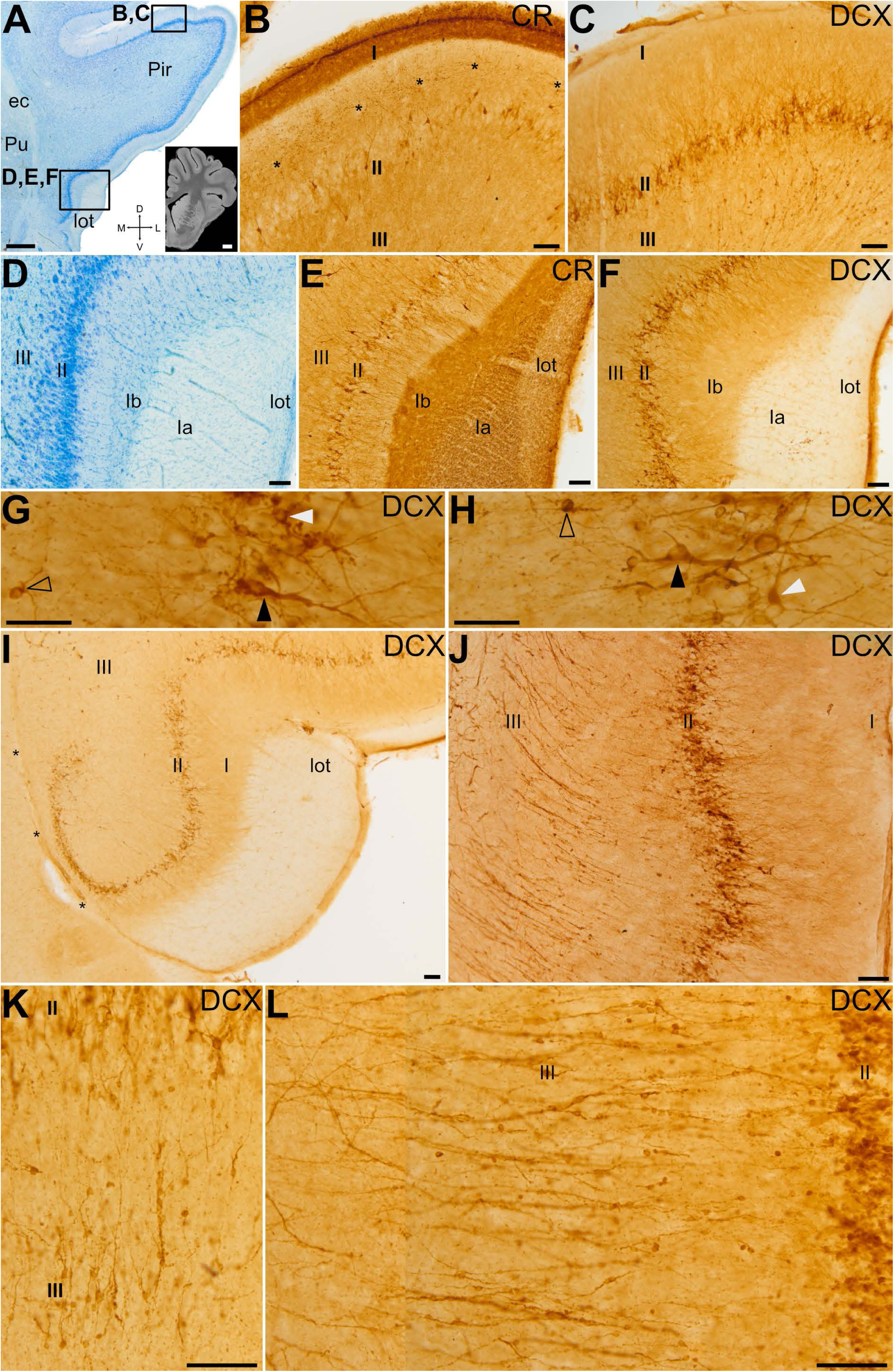
Distribution and migration of DCX+ cells in the piriform cortex. (A, D) Details of a Nissl stained frontal section at the anterior level of the piriform cortex (Pir); MRI image of a frontal section at a similar level (inset in A, taken from Saikali et al, 2010). Squared areas in A indicate the location of the higher magnification views shown in B-F. (B,C,E,F) Details of dorsal (B,C) and ventral (E,F) Pir immunostained for CR (B,E) or DCX (C,F). Note the CR+ lateral olfactory tract (lot) at the surface of Pir, serving as a landmark for its location. Asterisks in (B) point to CR+ fibers in Pir layer I parallel to the surface. Numerous DCX+ cells are located in cortical layer II at all levels, and display different morphologies, as seen in the details shown in G and H. Some have pyramidal (black arrowheads in G and H) or semilunar (white arrowheads in G and H) somas, with apical dendrites, while other cells are smaller and simpler (empty arrowheads in G and H). (I) Detail of ventral Pir, showing DCX+ cells arching inwards parallel to a blood vessel at the pallio-subpallial boundary (asterisks). (J) Detail of a posterior level of Pir, showing numerous DCX+ cells in layer II and cell chains radially traversing layer III. (K,L) Details of DCX+ cell chains in layer III of anterodorsal (K) and posteroventral Pir (L). For abbreviations, see list. Scales: A = 1mm; MRI A inset = 3mm; B-F, I-L = 100µm; G-H = 25µm

We also found numerous DCX+ cells distributed in layers II-III, with a very small round soma and short dendrites (empty arrowheads in Fig. 7G,H). In layer II, this cell type was preferentially located in the deep part. Furthermore, we observed other DCX+ cells with intermediate soma size and dendritic arborization. In the ventral part of Pir, near the boundary with the subpallium (marked by the presence of a radially oriented blood vessel, separating the Pir from the olfactory tubercle), small and medium-sized DCX+ cells arched inward and gradually changed from a compact to a looser organization (Fig. 7I).

In addition, in layer III we also detected some radially-oriented cellular chains, formed by a rosary of three or more DCX+ small somas, that traversed the Pir extending from the white matter deep to it (external and amygdalar capsules) to cortical layer II (Fig. 7J, details in K,L). While these migratory cell-like chains were seen at different levels of Pir, they were more defined and abundant at posterior levels (Fig. 7J, detail in L). They appeared to branch off from the external capsule, where similar chains of DCX+ cells were also observed and could be traced toward the vicinity of the Arc of the dorsal SVZ, following the lateroventral pallial migratory stream (Fig. 8). However, the internal capsule separated the observed chains from the SVZ. To better understand the possible origin and trajectory of the migratory cell-like chains, we analyzed them in sagittal and horizontal section planes (Fig. 8E-H). In these planes, we could observe a continuation at pre-callosal levels of the DCX+ cells exiting the SVZ (at Arc levels) with those of the external capsule (arrowheads in Fig. 8F).

**Figure 8.**
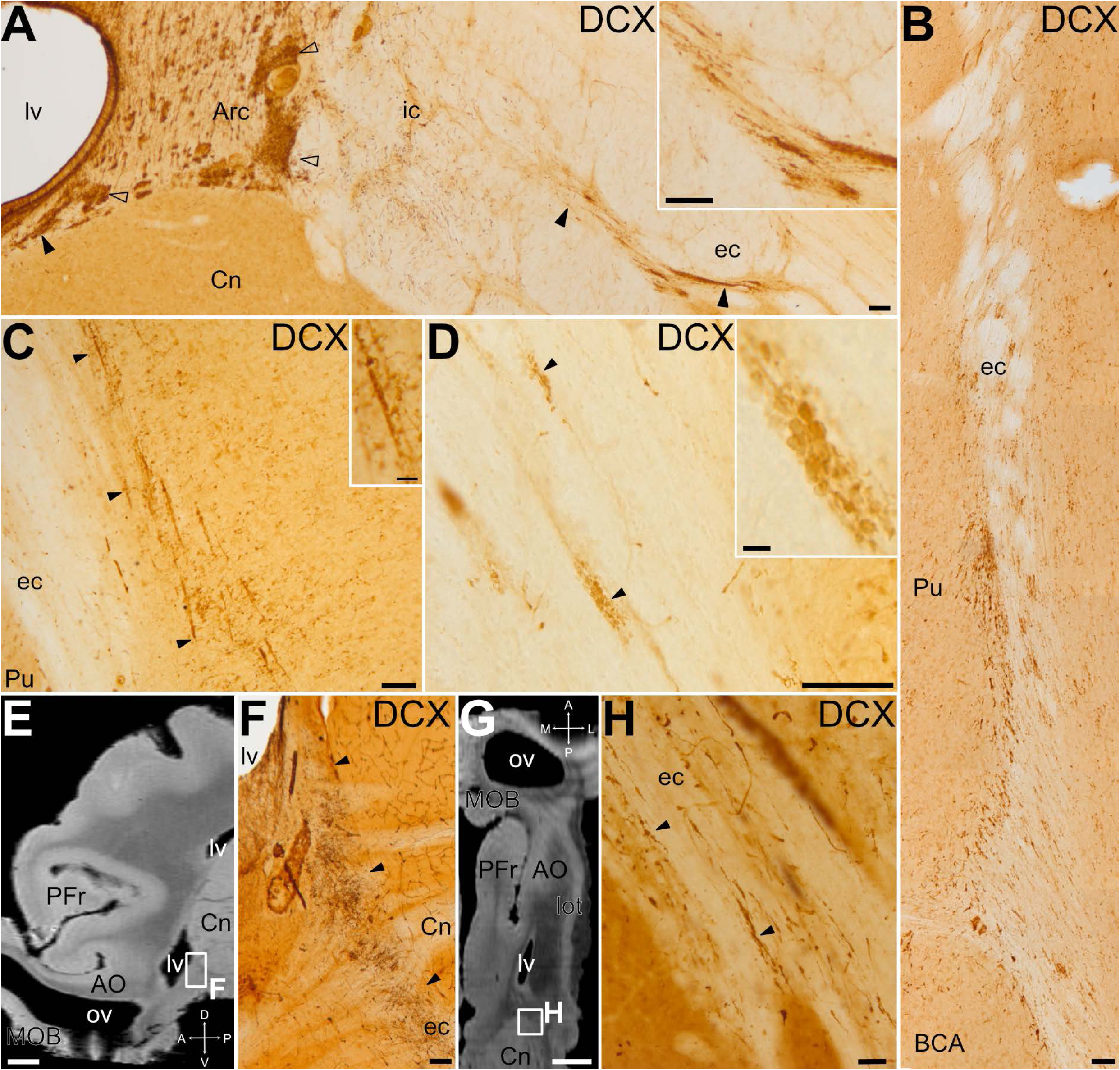
Distribution of DCX+ cell clusters and migratory-like chains through the external capsule. Details of coronal sections immunostained for DCX (A-D,F,H), showing DCX+ cell clusters in the dorsal SVZ/Arc (empty arrowheads) and through the external capsule (ec) (filled arrowheads), following the lateroventral pallial migratory stream (A, C and D from anterior levels, and B from a caudal level). Details of the DCX+ cell clusters are shown in A, C and D insets. In A, the internal capsule (ic) separates the SVZ/Arc from the DCX+ cell clusters observed more laterally in ec. In the lateral part of ec, DCX+ cells and chains were observed parallel to the fibers (arrowheads in C), and small clusters of DCX+ were also found intermingled between the fibers (arrowheads in D). (E,G) MRI images of sagittal (E) or horizontal (G) sections (taken from Saikali et al, 2010), at similar levels to the DCX immunostained images shown in F and H, respectively. (F) Detail of DCX+ clusters (arrowheads) connecting the SVZ (adjacent to the lateral ventricle, lv) and ec at a precallosal level. (H) Detail of clusters of small DCX+ cells along the ec (arrowheads). For abbreviations, see list. Scales: A-D, F = 100µm; H and inset in C = 25µm; inset in D = 10µm; E,G = 3mm

Regarding the phenotype of the DCX+ cells, double immunofluorescence labeling revealed DCX+/Ki-67+ cells within the chains, indicating that some of the migrating cells were undergoing division during migration (Fig. 9A). Moreover, we found other DCX+ cells in these migratory chains that expressed the transcription factor Brn2 (Fig. 9B).

**Figure 9.**
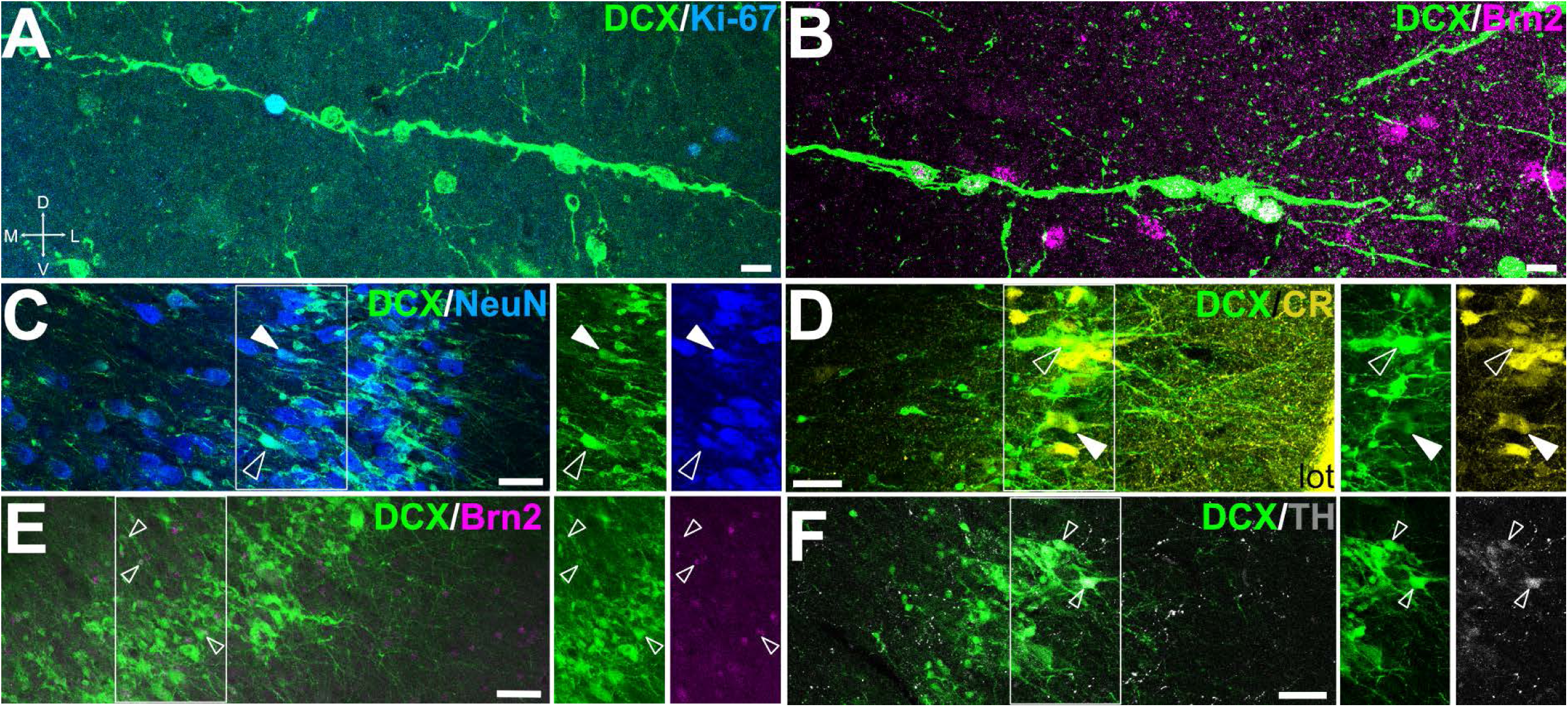
Proliferative capacity, migratory and mature phenotypes of DCX+ cells in the piriform cortex. (A-F) Confocal images of Pir details in frontal sections showing immunofluorescence for DCX (green) and either Ki-67 (blue in A), Brn2 (magenta in B, E), NeuN (blue in C), CR (yellow, D), or TH (gray in F). Squared areas in C-F are shown in separate channels on the right side of the merged image. (A) Some DCX+ cells in chains coexpress Ki-67, indicating that they continue proliferating while migrating. (B) DCX+ cells in chains show coexpression of Brn2. (C) DCX+ cells in layer II of the piriform cortex express the neuronal marker NeuN (arrowheads). Some DCX+ of layer II also express CR (arrowheads in D), Brn2 (arrowheads in E) or TH (arrowheads in F). In C and D, empty arrowheads show cells with high signal of DCX and low expression of NeuN or CR (suggesting an early maturation stage), while filled arrowheads point to cells with weak signal of DCX and high expression of NeuN or CR (putative more mature neurons). For abbreviations, see list. Scales: A, B = 10µm; C-F = 50µm

Almost all DCX+ cells in layer II showed coexpression with the neuronal marker NeuN (Fig. 9C). In addition, many of the DCX+ cells coexpressed CR (Fig. 9D) or Brn2 (Fig. 9E). In contrast, only a few of the DCX+ cells coexpressed TH (Fig. 9F). As in the OB, we observed differences in intensities of expression of the studied markers, with opposing gradients: cells with strong expression of DCX and low expression of NeuN or CR (empty arrowheads in Fig. 9C,D), and cells with low expression of DCX and high expression of NeuN or CR (filled arrowheads in Fig. 9C,D), which could be linked to differences in the maturity degree of the cells.

### DCX expression in cortical amygdalar areas and entorhinal cortex

Caudally to the Pir, we studied the presence of DCX+ cells in two other secondary olfactory areas: the cortical amygdalar areas (Co) and the entorhinal cortex (EC) (Martínez-Marcos, 2009; Shipley et al, 2008). DCX+ cells were found in both Co and EC; however, in contrast to the Pir, they exhibited a patchy distribution (Fig. 10A and D, details in B and E).

**Figure 10.**
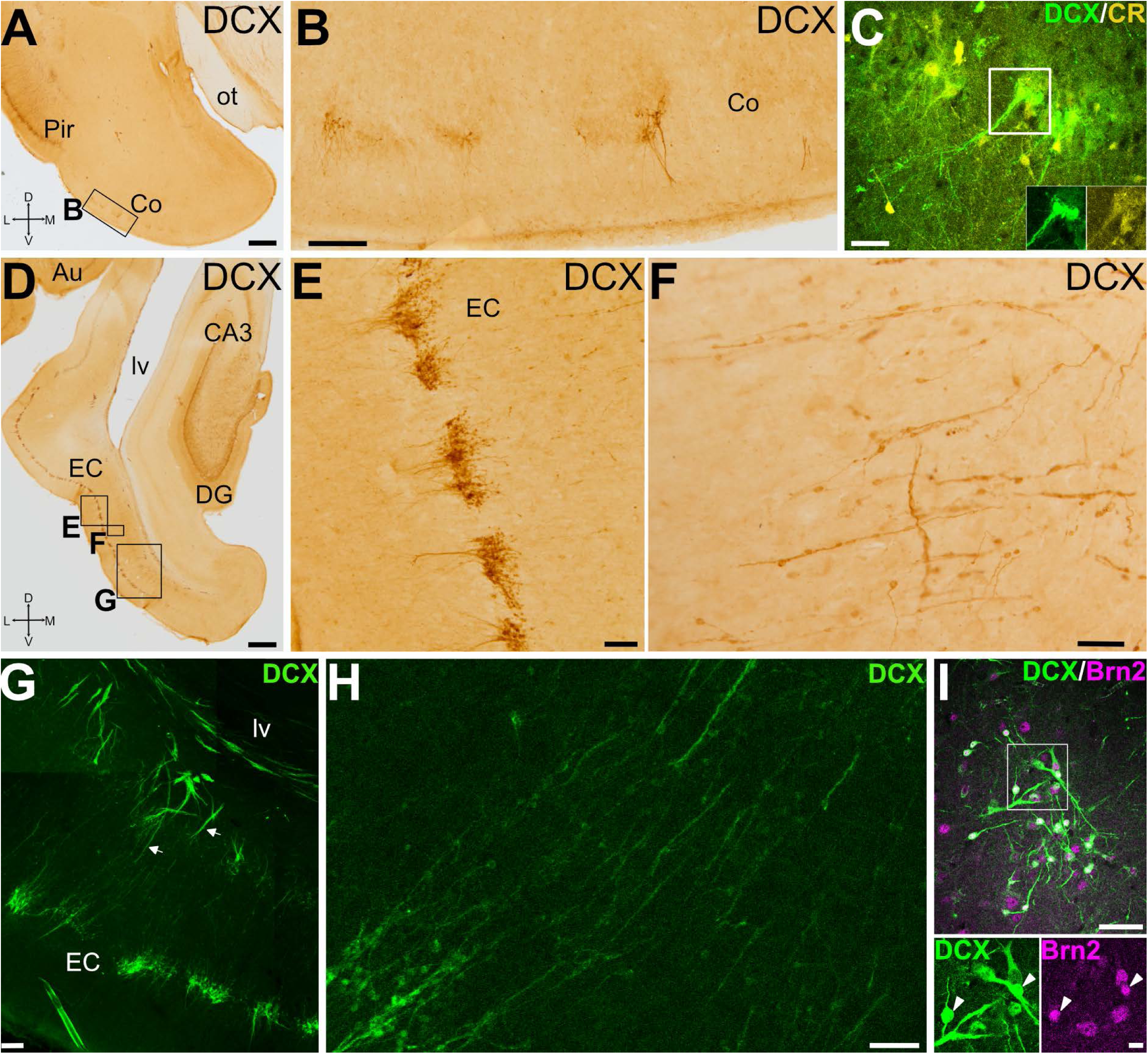
DCX expression in cortical amygdalar area and entorhinal cortex. (A,B) Frontal sections at the level of the cortical amygdala (Co) immunohistochemically stained for DCX. Squared area in A is shown at higher magnification in B. Few DCX+ cells are grouped in small patches or clusters through Co, and some of the cells display a distinct apical dendrite. (C) Confocal image of a Co detail in a frontal section, showing double immunofluorescence for DCX (green) and CR (yellow). Squared area in C is shown with split channels in insets. (D-F) Frontal sections at the level of the entorhinal cortex (EC) immunostained for DCX. Squared areas in D are shown at higher magnification in E and F. (E) DCX+ cells are grouped in small patches or clusters through the EC layer II. (F) In the inner part of the EC, DCX+ cells form radially-oriented migratory-like chains. (G) Confocal image of EC (level like the G squared area in D) showing DCX immunofluorescence. DCX+ cell chains (arrows) depart from the SVZ around the ventro-caudal part (temporal horn) of the lateral ventricle (lv). (H) Detail from G showing the cell chains in the deep part of EC. (I) Confocal image of details of layer II EC cells, showing double immunofluorescence for DCX (green) and Brn2 (magenta). Squared area in the merged image is shown at higher magnification in separate channels below. Many DCX+ cells in the clusters co-express Brn2 (arrowheads). For abbreviations, see list. Scales: A, D = 1mm; B, E, G = 100µm; C, F, H, I = 50µm; insets with split channels in I = 10µm

In the Co, DCX+ cells were observed mainly in its anterior part. Their somas were lightly immunoreactive and formed patches in layer II (Fig. 10B). Most of these cells had small-rounded somas, and some of them had a prominent dendritic process oriented radially toward the surface (Fig. 10B). This area was rich in CR+ neuropile and cell bodies, and some of the cells coexpressed DCX (Fig. 10C).

Regarding the EC, clusters of intensely immunoreactive DCX+ cells were found in layer II (Fig. 10D), mainly in lateral EC. These clusters were composed of numerous DCX+ cells with small-rounded somas and remarkable dendritic processes oriented toward the surface, which showed a tendency to converge with those of other DCX+ cells (Fig. 10E). We also observed thin cellular chains formed by three or more DCX+ small somas, which traversed the EC deep layers, departing from similar migratory chains in SVZ parallel to the temporal horn of the lateral ventricle and, more laterally, in the white matter carrying the connection fibers between EC and the lateroventral pallium (Fig. 10F-H). The majority of DCX+ somas in EC coexpressed the glutamatergic transcription factor Brn2 (Fig. 10I).

## DISCUSSION

### Immature cells in the swine olfactory system at juvenile stages

Our results show the existence of numerous immature cells expressing DCX throughout the primary and associative structures related to olfaction in the brain of juvenile swine. Previous studies showed the presence of DCX+ cells in the OB and RMS of piglets (baby pigs until weaning, which occurs at about 21 days) (Costine et al. 2015; Porter et al. 2022), but our finding of abundant immature cells in 2.5 to 3.5 months-old animals suggests that there is still a great level of neural plasticity in the olfactory system at prepuberal postnatal stages. This agrees with studies in other artiodactyl mammals, such as the sheep (Piumatti et al. 2018), as well as with findings in other large-brained mammals with a relatively prolonged postnatal development, such as human (Paredes et al., 2016; Sorrells et al., 2019) and non-human primates (Bernier et al. 2002; La Rosa et al. 2020). The abundance of immature cells, many with a neuronal profile based on expression of NeuN or other neuronal markers, is much larger and spread over more cortical and non-cortical pallial areas in these large brained gyrencephalic mammals compared to small brained lissencephalic mammals, such as murine rodents (Bonfanti and Nacher 2012; La Rosa et al. 2020; Bonfanti and Seki 2021). It has been suggested that such immature neurons might represent a reservoir of undifferentiated cells that could contribute to the postnatal plasticity of pallial areas in large-brained mammals with high cognitive capacities (La Rosa et al. 2020).

Indeed, brain plasticity is critical to allow adaptive changes at functional and structural levels (Bonfanti, 2006), and may be achieved by adding new neurons (neurogenesis) and/or by modifying synaptic connections (synaptogenesis). However, postnatal neurogenesis is limited in terms of quantity of neurons produced, location of neurogenic niches and variety of cellular phenotypes produced and shows a high variability between species (reviewed in Gould 2007; Barnea and Pravosudov 2011; Paredes et al. 2015; La Rosa et al. 2020; Zupanc 2021). In mammals, adult neurogenic niches have only been consistently found in the SVZ and RMS, producing new neurons for the OB, and in the subgranular zone (SGZ) of the dentate gyrus in the hippocampal formation (Lledó et al. 2006; Gould 2007). In humans, the capacity of producing new neurons appears to dramatically drop after early infancy (Paredes et al. 2015; Sorrells et al. 2018). Based on the absence of mitotic markers in DCX+ cells (Brus et al. 2013; Piumatti et al. 2018; La Rosa et al. 2020), it has been suggested that the immature cells found throughout many cortical/pallial areas in large-brained mammals are not born postnatally, but prenatally, and remain in quiescent immature form until needed, thus providing a non-neurogenic type of plasticity that is more prevalent in large-brained mammals (Bonfanti et al. 2023). However, in juvenile swine we found many chains of cells with small soma and migratory morphology (as described by Rousselot et al. 1995; Lois et al. 1996) in the SVZ, RMS, external and amygdalar capsules adjacent to piriform cortex and white matter related to entorhinal cortex, as well as in all olfactory cortical areas from rostral to caudal levels. Moreover, in spite of the short period of overlap between DCX and Ki-67 expression (Fig. 1), we also found many cases of DCX+ cells that coexpressed Ki-67 not only in SVZ and RMS, but also in the migratory chains of several cortical olfactory areas, such as AO and Pir, in agreement with the findings of Porter et al. (2022) in piglet. Some DCX+ cells with similar migratory morphology and position expressed Brn2, a transcription factor involved in the maturation of different subpopulations of glutamatergic neurons in both the cortex (Dominguez et al. 2012) and the cerebellum (Wu et al. 2022; Casoni et al. 2024). Thus, although we cannot discard that most immature cells found postnatally may represent non-newly generated quiescent cells, the present results suggest that new neurons are also produced in several olfactory pallial areas of the juvenile swine. Nevertheless, there might be a drop in neurogenesis after puberty in swine, and in adults cell proliferation in the cortex might be restricted to glial cells, which would agree with observations in sheep, humans and other mammals (Brus et al. 2013; Piumatti et al. 2018; Sorrells et al. 2019; La Rosa et al. 2020). However, one study in two species of adult monkeys showed production of new neurons that incorporated in the Pir and parts of the pallial amygdala after injections of the proliferation cell marker BrdU (Bernier et al. 2002). It is unclear whether the discrepancy between the findings of this article and those in humans and other mammals might be due to species differences or to technical issues. For example, Bernier et al. (2002) carried out intraventricular injections of BrdU (doses: 50mg/kg) in adult monkeys twice a day for 3 days and sacrificed animals at several times (up to 21 or 28 days, depending on the monkey species) after the last injection, observing larger numbers of new neurons in the pallial mantle following longer periods of survival. In contrast, Brus et al. (2013) and Piumatti et al. (2018) (La Rosa et al. 2020, referred to these two publications) studied the existence of adult neurogenesis in pregnant ewe that received four intravenous injections of BrdU (doses: 20mg/kg), waiting between 30 and 120 days to sacrifice, and concluded that DCX+ cells were generated before birth. These differences in doses and delivery methods could have an impact on the availability of BrdU to brain dividing cells. Additional studies would be required to investigate these possibilities.

### Phenotype of the DCX+ cells of the OB and AO, and their possible origins and migratory pathways

Our data showed the presence of DCX+ cells in the olfactory bulbs (mainly in MOB) of juvenile swine, most of which express NeuN, indicating a neuronal phenotype. In the OB of rodents, non-human and human primates, most newly generated cells become granular interneurons (94%) and periglomerular cells in a smaller proportion (4%) (Lledó et al. 2006). These new cells acquire a GABAergic phenotype and express CB, CR or TH (Kohwi et al. 2005, 2007). However, our results in swine OB showed that DCX+ neurons were mostly located in the GCL, but extremely few cells were also observed in the GL. We found some TH+ periglomerular cells in the GL, as in other mammalian species (Bagley et al. 2007; Panzanelli et al. 2007; Angelova et al. 2018), but we could not detect coexpression with DCX. One possible explanation of this discrepancy is that in swine OB the levels of neurogenesis of periglomerular cells could be too low making its detection difficult. It is also possible that the temporal pattern of differentiation of dopaminergic (TH+) periglomerular cells do not overlap the transitory expression of DCX.

In the GCL of juvenile swine, in addition to NeuN, we found many cases of DCX+ cells coexpressing the calcium-binding protein calretinin, in line with results in OB of other mammals (Kohwi et al. 2005, 2007). Interestingly, these CR+ cells extend processes radially oriented towards mitral cells, and might establish contacts with them, in agreement with previous studies in mouse demonstrating that these cells establish dendrodendritic contacts with lateral dendrites of mitral cells in mouse (Takahashi et al. 2018). In rodents, newly generated CR+ interneurons of GLC integrate in a network for modulation of mitral cells and their transmission of olfactory information (Takahashi et al. 2018) and this network appears to be highly plastic, based on their continuous renewal and expression of DCX. Studies in mice have shown that adult-generated CR+ cells of GLC are involved in complex odor discrimination (Hardy et al. 2018).

In mice, a subpopulation of adult-generated glutamatergic periglomerular neurons has also been described in the OB, which appears to originate from Neurog2 and Tbr2 progenitors of the dorsal part of SVZ (Brill et al. 2009). In our swine samples, Brn2 expression was mainly observed in EPL (possibly tufted cells) and mitral cells as previously described in other mammals (Hagino-Yamagishi et al. 1999). Mitral cells of swine OB also coexpressed CR, as expected based on previous results in other mammals (Wouterlood and Härtig 1995; Hagino-Yamagishi et al. 1999), but they did not express DCX. However, we found abundant cases of DCX+ neuroblasts coexpressing the glutamatergic transcription factor Brn2 in the RMS. The fate of these immature neurons is still unclear, but it is likely that many of them eventually integrate into different subregions of the AO, as discussed below.

In several vertebrate species, newly-generated neurons of the OB are known to originate from a subpopulation of stem cells located in the subventricular zone of the lateral ventricle (SVZ) and migrate to the OB through the RMS, where some cells maintain the proliferative capacity (Pérez-Cañellas and García-Verdugo 1996; Font et al. 2001; Merkle et al. 2007; Kelsch et al. 2007; Young et al. 2007; Gil-Perotin et al. 2009; Sawamoto et al. 2011; Malik et al., 2012; Ponti et al. 2013; Paredes et al. 2016). Our results in swine agree with these findings. Using the proliferation marker Ki-67, we identified abundant dividing cells in the SVZ, the RMS, and the periventricular region surrounding the rostral extension of the lateral ventricle (or olfactory ventricle) in juvenile swine brains. Some of them coexpressed DCX and had a migratory neuroblast morphology, suggesting that they were dividing during migration, in agreement with previous studies in different mammals, including piglets (Costine et al. 2015; Porter et al. 2022) and juvenile swine (Torrijos-Saiz et al. 2025). These DCX+ cells in the outer part of SVZ and in RMS showed an elongated morphology, which agrees with their ultrastructural features of migratory neuron, with reduced cytoplasm and typical microtubule organization oriented parallel to the leading process (Torrijos-Saiz et al. 2025). While many of the proliferating cells appear to be involved in production of new neurons, we cannot discard that some of the DCX+/Ki-67 cells in SVZ will acquire an oligodendrocyte fate, as found in some studies (Boulanger and Messier 2017).

The swine SVZ probably contains discrete niches, analogous to those described in rodents (Batista-Brito et al. 2008; Fernández et al. 2011; Merkle et al. 2007; Weinandy et al. 2011; Kohwi et al. 2007; Young et al. 2007). In rodents, each niche has a unique molecular profile and follows its own spatiotemporal pattern to produce specific olfactory interneurons subtypes (Batista-Brito et al. 2008; Fernández et al. 2011; Merkle et al. 2007; Weinandy et al. 2011; Kohwi et al. 2007; Young et al. 2007). Further research is needed to confirm this hypothesis, but there is evidence in piglets that DCX+ migratory neuroblasts in the SVZ clusters express GABA, and some of them contain CR, PV or NPY (Porter et al. 2022). As noted above, we also found DCX+ cells coexpressing CR in the GCL of the MOB of juvenile swine. At least in mouse, only CR, CB and TH interneurons undergo postnatal/adult renewal, but not PV interneurons (Young et al. 2007). If this is similar in swine, the cases of DCX+/PV+ cells found in piglets may represent a non-neurogenic type of plasticity.

The swine SVZ that produces the DCX+ neuroblasts of the RMS is mostly located above the striatum (caudate nucleus) and relates to an accumulation of DCX+ cell clusters previously referred as Arc (Torrijos-Saiz et al. 2025; see also Paredes et al. 2016 in the infant human). In piglets, the cell clusters of this sector express the transcription factor Sp8 (Porter et al. 2022), which in mouse is associated to the dorsal/caudal lateral ganglionic eminence during development (Ma et al. 2012) and to the RMS and the development of OB interneurons (Li et al. 2018). In humans, in addition to the RMS, another branch of migratory neuroblasts split off the Arc and produce interneurons for the prefrontal cortex, but both migratory routes are only observed in early infancy and dramatically drop after (Paredes et al. 2016). The prefrontal migratory route also appears to be present in other gyrencephalic mammals (Ellis et al. 2018; Pencea et al. 2001), including piglets (Porter et al. 2022). However, the dorsal part of the postnatal SVZ associated to the Arc is complex. Studies in mouse showed that it contains progenitors expressing pallial (such as Emx1) and subpallial (such as Dlx5/6 and Gsh2) transcription factors (Kohwi et al., 2007; Young et al., 2007). Moreover, a subset of Emx1-lineage cells in this SVZ region as well as their progeny in the OB express the transcription factor Dlx2, which acts upstream of Dlx5/6 (Kohwi et al., 2007). Most adult-produced OB interneurons (70%) derive from subpallial SVZ progenitors, but it appears that Emx1 progenitors also produce part of the OB GABAergic interneurons, including some of the CR+ interneurons (Kohwi et al. 2007; Young et al. 2007).

In addition, the SVZ may also include pallial progenitors that produce glutamatergic cells for different pallial sectors. Around the ov, in relation to the AO, we found many DCX+ cells coexpressing the transcription factor Brn2. It is thus possible that the AO receives GABAergic interneurons as well as glutamatergic neurons through the RMS. Moreover, we found cases of DCX+/CR+/Brn2+ triple-labeled cells in the AO. Thus, a subpopulation of AO CR immature cells may be glutamatergic although this would require confirmation using additional markers, such as Tbr1 (Brunjes and Osterberg 2015). The incorporation of newly generated neurons, both GABAergic and glutamatergic, may contribute to maintaining functional plasticity within this area, which is very large and complex in swine (Brunjes et al. 2016). One possible mechanism for modulating this plasticity involves olfactory sensory inputs (Lledó et al. 2006; Bastien-Dionne et al. 2010). In this regard, it is noteworthy that we observed axons branching off from the lot toward the DCX+ cells located in the AO.

As described in rodents and primates (Alvarez-Buylla and García-Verdugo 2002; Gil-Perotin et al. 2009), in juvenile swine the DCX+ migratory cells of the RMS form aggregates and appear to move along each other forming chains, as also suggested by electron microcopy data showing frequent adherens junctions between neuroblasts (Torrijos-Saiz et al. 2025). This strategy is similar to the migration described during embryonic development and allows these neuroblasts to complete the long pathway to the OB while dividing and maturing (Alvarez-Buylla and García-Verdugo 2002). However, the migratory capacity of these immature cells in swine needs to be confirmed using migratory assays in organotypic slice cultures or in vivo.

We also observed that the chains of DCX+/Ki-67+ migratory cells of the RMS were surrounded by TH+ fibers, which were parallel to DCX+ cell chains, suggesting that they might play a role as a guidance for these migratory-like cells. These data are aligned with results in mice that showed that dopaminergic TH-denervation decreases neuroblast proliferation and migration to the OB (Höglinger et al. 2004). In addition, we observed that some of the DCX+/Ki-67+ migratory cells were parallel and adjacent to blood vessels, in agreement with a similar finding in piglets (Porter et al. 2022). This apparent association suggests that blood vessels could also contribute to guide and perhaps modulate the migration of these cells during the postnatal period, a possibility that requires further investigation.

### Phenotype of DCX+ cells in Pir and other olfactory cortical areas and their possible origin(s) and migratory pathways

Our data in juvenile swine also showed the presence of DCX+ cells in additional olfactory areas, including the piriform (Pir), amygdalar (Co) and entorhinal cortices (EC). In these areas, many DCX+ were located in layer II and expressed NeuN.

In the Pir of juvenile swine, some DCX+ cells showed a mature morphology (pyramidal or semilunar), with larger cell bodies and apical dendritic branches. Many of them also contained Brn2 (i.e. they may be glutamatergic) and some also were CR+. Similar cell subtypes have been described in the Pir layer II of different mammals, as well as in layer II of neocortex and entorhinal cortex, and it has been suggested that they may represent different stages in the maturation of pyramidal neurons (Bonfanti and Nacher 2012; Piumatti et al. 2018). This resembles the findings in the sheep and other mammals, where some of the DCX+ neurons of Pir layer II exhibit pyramidal-like and semilunar morphologies (Bonfati and Nacher 2012; Rubio et al. 2016) and many display a glutamatergic phenotype, based on their expression of the transcription factor Tbr1 (Luzzati et al. 2009; Piumatti et al. 2018). While DCX+ glutamatergic neurons likely originate form pallial progenitors, only a few DCX+ neuroblasts of the olfactory cortical areas appear to be subpallial-derived interneurons, based on their expression of Sp8 (sheep; Piumatti et al. 2018). In the Pir of juvenile swine, we observed a subset of DCX+ cells with expression of TH, which may represent a different interneuron subtype. Moreover, we identified smaller DCX+ cells, possibly less mature, that did not express NeuN. Some of these small cells could be glial cells as explained below.

In juvenile swine we found chains of small DCX+ migratory-like cells that split off the external/amygdalar capsule and radially traversed deep layers of Pir. Similar chains of DCX+ migratory-like cells were found in the white matter deep to EC and traversing the deep layers of this cortical area. Some DCX+ cells in the migratory chains expressed Ki-67, and many expressed Brn2. Thus, it appears that cells of these chains continue proliferating while migrating and, at least some of them, might be producing new glutamatergic neurons for the Pir and EC, which appear to mature and integrate into layer II. In addition, we cannot discard that some small DCX+ cells are immature oligodendrocytes, similar to those described in swine EC by Liu et al. (2021). Our results agree with the finding of newborn neurons in Pir and pallial amygdala of adult monkeys after injection of BrdU (Bernier et al. 2002). These authors showed that the new cells are produced in a neurogenic niche of the SVZ near the temporal horn of the lateral ventricle and, from there, migrate along the temporal migratory stream, in parallel to the fibers of the external capsule, to reach their destination. In our study, we found groups of DCX+ migratory-like cells in the external and amygdalar capsules, an observation that has also been made in sheeps (Piumatti et al. 2018). Moreover, in swine the chains of DCX+/Ki-67 migratory cells located in the deeper layers of the Pir appeared to be continuous with, and possibly emerging from the capsule. To try to understand the origin of those cells, we analyzed brain sections in different planes and found that, at precallosal levels, the SVZ has a lateroventral branch of DCX+ cells that enter the external capsule. This could be a way of distribution, but – as mentioned above – our results showed that proliferation continues in the migratory cell chains, as it happens in the RMS. However, it is likely that additional niches exist in the SVZ adjacent to posterior/temporal levels of the lateral ventricle at least in some gyrencephalic mammals (Bédard and Parent 2004; Bernier et al. 2002; Cai et al. 2009; Chawana et al. 2013; Ellis et al. 2018; Marlatt et al. 2011; Tonchev et al. 2003), a possibility that was also suggested in piglets (Costine et al. 2015) and which requires further investigation.

Overall, these findings suggest that the proliferative niches described in the SVZ adjacent to the rostral and dorsal regions of the lateral ventricle are quite complex and appear to extend along the rostro-caudal axis, giving rise to different migratory streams and neuronal phenotypes, including both GABAergic and glutamatergic neurons. In addition, these proliferative niches may also give rise to glial cells, some of which may express DCX+ (based on coexpression with oligodendrocyte precursor markers; Boulanger and Messier 2017).

Contrary to the postnatal neurogenesis in the SVZ/OB and in the SGZ/hippocampus, which have been demonstrated throughout vertebrates, the production of new neurons, born after birth, that incorporate to other cortical structures throughout life remains controversial (Feliciano et al. 2015). In fact, the finding of immature neurons expressing DCX+ and/or PSA-NCAM in different cortices, such as the Pir, the EC and different parts of the pallial amygdala is interpreted by many authors as a form of non-neurogenic plasticity (reviewed by La Rosa et al. 2020; Bonfanti et al. 2023; Ghibaudi et al. 2023a,b, 2025). However, while we agree that DCX+ cells in layer II are not proliferating, in the juvenile swine we found evidence suggesting that new neurons, mostly Brn2+ and possibly glutamatergic, can be postnatally added to several of these areas. Thus, neurogenic and non-neurogenic plasticity may exist in Pir and perhaps other olfactory cortical areas of swine, at least until puberty.

### Functions of immature neurons in the olfactory system

Olfactory structures are essential for maintaining individual responses to familiar contexts, while preserving the capacity to respond to novel conditions (Bernier et al. 2002; Carleton et al. 2003; Lledo et al. 2006). In farms, when the weaning process ends, juvenile swine are separated from their mothers and caged in new groups. These ongoing modifications at environmental and social level, taken together with physiological changes to achieve sexual maturity, might require numerous behavioral adaptations to maximize their survival chances. Olfaction in swine is crucial to pheromone detection and sexual interaction, to recognize the members of the social group and establish dominance-submission roles between them (Dorries et al. 1995; Kristensen et al. 2001; McGlone 1985; McGlone et al. 1987; Sink 1967). Considering this, it is not surprising that olfactory structures of these puberal animals show such an impressive number of immature neurons.

Postnatal neurogenesis adds and/or replaces neurons according to surrounding requirements to allow a better adaptation to life and, not surprisingly, it seems to be well-preserved throughout invertebrates and vertebrates (Barnea and Pravosudov 2011; La Rosa et al. 2019). The role of newly generated neurons incorporated in the OB of different species remains unclear. However, research done in rodents combining lesion approaches, electrophysiological designs and permanent labeling techniques (BrdU and/or ^3^H-thymidine) reported that the survival and integration of adult-generated granular neurons in the OB is determined in a sensory experience-dependent manner (Yamaguchi and Mori 2005). Moreover, some authors suggested that new-incorporated cells in the GCL maximize differences in odor representations, to improve olfactory discrimination and olfactory memory (Carleton et al. 2003; Lledo et al. 2006; Petreanu and Alvarez-Buylla 2002; Lledo and Valley 2016; Hardy et al. 2018). The functional explanation of postnatal-generated neurons in cortical olfactory structures is less explored. Nevertheless, some authors consider this process parallel with the addition of new neurons in the OB (Bernier et al. 2002).Although a fraction of these immature neurons might be generated postnatally, the large number of DCX cells found in different pallial areas suggests that many of them might remain in quiescent form and can later mature and be recruited into functional circuits when physiologically required (Palazzo et al. 2018; Sorrells et al. 2019; La Rosa et al. 2020; Bonfanti and Couillard-Després 2023).

Overall, our results highlight the swine as a very valuable model to study brain plasticity throughout ontogenesis. Unlike murine rodents, the swine has a relatively large, gyrencephalic brain with anatomical and developmental features more similar to those of humans (Kostović et al. 2019; Liu et al. 2021), making it a better model to extrapolate findings on the role of postnatal plasticity and their potential for regeneration.

## Acknowledgements

We deeply thank all Agencies that funded our research: the Spanish Ministerio de Ciencia, Innovación y Universidades and Agencia Estatal de Investigación, MICIU/AEI/10.13039/501100011033 and FEDER-EU (Grant no. PID2023-151927OB-I00 to LM and ED), and the AGAUR/Generalitat de Catalunya (2021 SGR 01359 to LM). We also thank the technicians and other staff of the Department of Experimental Medicine, the Service of Proteomics and Genomics of the University of Lleida, as well as the veterinarians, technicians and other staff of the Center for Experimental Research in Applied Biomedicine (CREBA) of the Institute for Biomedical Research of Lleida (IRBLleida).

**Supplementary figure Negative controls for immunohistochemistry and immunofluorescence**

Horizontal (A-C, G-J’’) and sagittal (D-F, K-K’’) sections at the level of anterior olfactory area (AO), piriform cortex (Pir), entorhinal cortex (EC), and main olfactory bulb (MOB), processed for immunohistochemistry (A-I) or triple immunofluorescence (J-K’’) omitting primary antibodies. No specific immunoreactivity was detected, although the anti-goat secondary antibodies produced some background (without clear cell labeling), possibly due to the phylogenetic proximity between goat and swine (G-I, J’). In addition, blood vessels were labeled with some of the secondary antibodies (C, G, I, J, J’’, K-K’’). Mediolateral and dorsoventral axes are indicated in A for orientation. For abbreviations, see list. Scales: A = 500µm (applies to A-I); J = 100µm (applies to J-K’’).

## ABBREVIATIONS

A: anterior
AO: anterior olfactory area
AOal: anterior olfactory area, anterolateral part
AOam: anterior olfactory area, anteromedial part
AOd: anterior olfactory area, dorsal part
AOl: anterior olfactory area, lateral part
AOm: anterior olfactory area, medial part
AOpl: anterior olfactory area, posterolateral part
AOpm: anterior olfactory area, posteromedial part
AOB: accessory olfactory bulb
Au: auditory cortex
BCA: basal complex of the amygdala
BrdU: bromodeoxyuridine
BST: bed nucleus of the stria terminalis
CA3: cornu Ammonis 3
CB: calbindin
cc: corpus callosum
Cerb: cerebellum
Cg: cingulum
Cl: claustrum
Cn: caudate nucleus
Co: cortical amygdalar area
cp: cerebral peduncle
CR: calretinin
D: dorsal
dAO: dorsal part of the anterior olfactory area
DCX: doublecortin
DG: dentate gyrus
Dlx2: distal-less homeobox 2 transcription factor
Dlx5-6: distal-less homeobox 5-6 transcription factors
DMSO: dimethyl sulfoxide
EC: entorhinal cortex
ec: external capsule
Emx1: empty spiracles homeobox 1 transcription factor
EPL: external plexiform layer
fx: fornix
GABA: gamma aminobutyric acid
GCL: granule cell layer
GL: glomerular layer
GP: globus pallidus
Gsh2 (Gsx2): genetic-screened homeobox 2 transcription factor
HF: hippocampal formation
IC: insular cortex ic internal capsule
IV: intravenous
L: lateral
lot: lateral olfactory tract
lv: lateral ventricle
M: medial
M1: primary motor cortex
MCL: mitral cell layer
Me: medial amygdala
MOB: main olfactory bulb
MRI: magnetic resonance imaging
NeuN: neuronal nuclear protein
Neurog2: neurogenin 2
NPY: neuropeptide Y
OB: olfactory bulb
och: optic chiasm
ot: optic tract
ov: olfactory ventricle
P: posterior
PB: phosphate buffer
PB-Tx: phosphate buffer containing Triton-X 100
PFA: paraformaldehyde
PFr: prefrontal cortex
Pir: piriform cortex
pM: premotor cortex
PRh: perirhinal cortex
PSA-NCAM: Polysialylated-neural cell adhesion molecule
Pu: putamen
PV: parvalbumin
RMS: rostral migratory stream
Rt: reticular prethalamic nucleus
S1: primary somatosensory cortex
Sa: somatosensory association cortex
Se: septum
SGZ: subgranular zone of the dentate gyrus
SVZ: subventricular zone
Tbr1: T-box brain transcription factor 1
Tbr2 (Eomes): T-box brain transcription factor 2
Th: thalamus
TH: tyrosine hydroxylase
TMS: temporal migratory stream
Tu: olfactory tubercle
V: ventral
V1: primary visual cortex
VZ: ventricular zone

## STATEMENTS & DECLARATIONS

## Author Contributions

This work is part of the Doctoral Thesis of JF, supervised by ED and LM. JF contributed to the experimental design, conducted immunohistochemical and immunofluorescence experiments, analyzed data, prepared the figures and wrote the first draft of the manuscript. FESA-R performed Nissl and immunohistochemical staining and participated in the analysis of the material. RN performed immunohistochemical and immunofluorescence staining and participated in the analysis. ED, LM and JF provided the study samples. LM and ED conceived and designed the study, supervised the project, secured funding, contributed to samples characterization, interpretation of results, and improving the manuscript. All authors approved the final manuscript and agreed with the results presented.

## Data Availability

The most relevant data are included in the Figures of this article. Additional data is available upon request and agreement.

## Ethics Approval

This study was performed in line with the 3Rs principles (Russell and Burch, 1959). Brain samples were obtained from animals that were previously used for practicing and improving surgical procedures by M.D. surgeons of the Arnau de Vilanova University Hospital of Lleida. Since brains were extracted once animals were euthanized, Animal Research Ethics Committee of the IRBLleida confirmed that no ethical approval is required.

