## Supplementary figures and images for "Postnatal plasticity in the olfactory system of the juvenile swine brain"

### Supplementary Figure 1

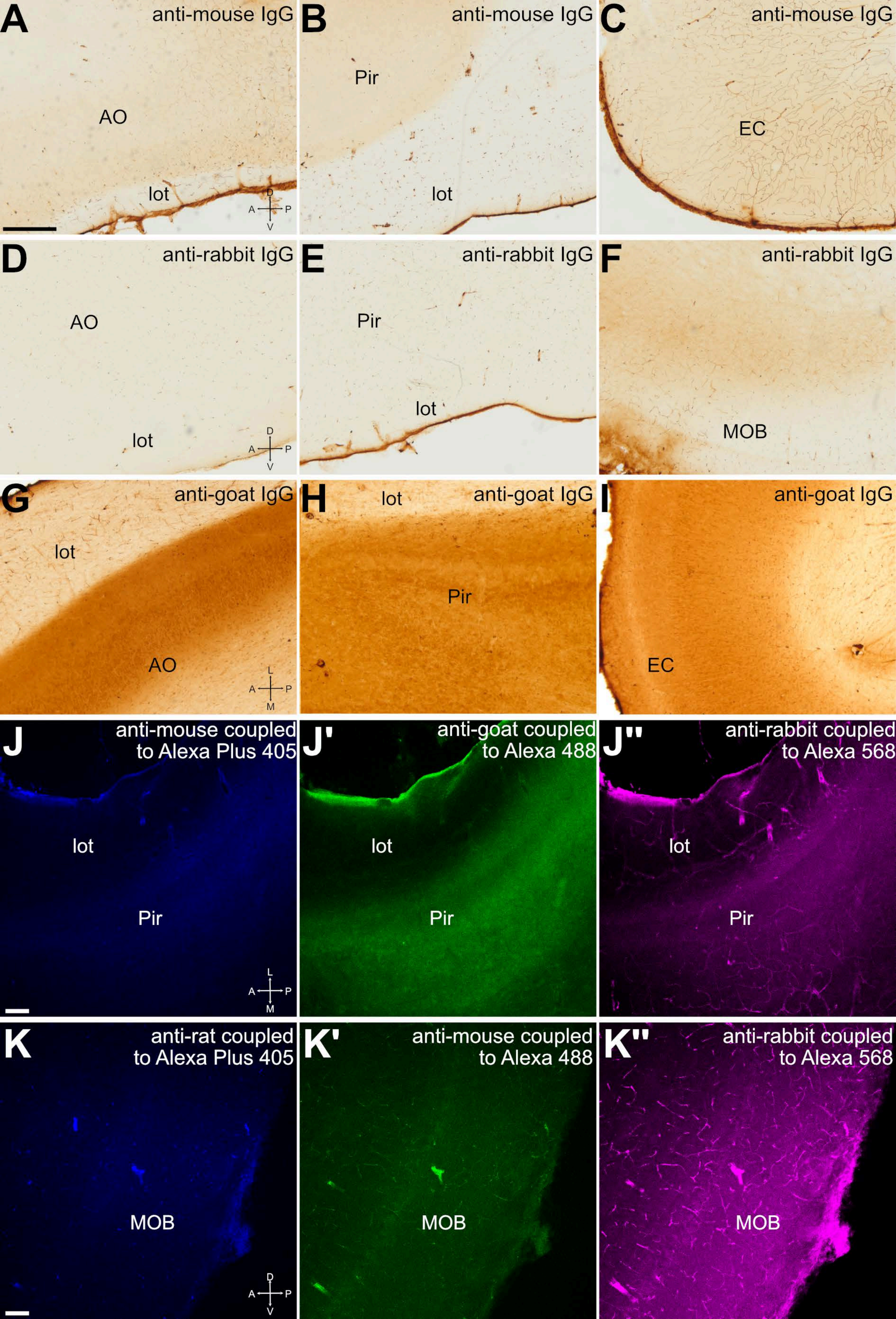
